# Tissue fixation effects on human retinal lipid analysis by MALDI imaging and LC-MS/MS technologies

**DOI:** 10.1101/2021.04.29.442044

**Authors:** Ankita Kotnala, David M.G. Anderson, Nathan Heath Patterson, Lee S. Cantrell, Jeffrey D. Messinger, Christine A. Curcio, Kevin L. Schey

## Abstract

Imaging mass spectrometry (IMS) allows the location and abundance of lipids to be mapped across tissue sections of human retina. For reproducible and accurate information, sample preparation methods need to be optimized. Paraformaldehyde fixation of a delicate multilayer structure like human retina facilitates the preservation of tissue morphology by forming methylene bridge cross-links between formaldehyde and amine/ thiols in biomolecules; however, retina sections analyzed by IMS are typically fresh-frozen. To determine if clinically significant inferences could be reliably based on fixed tissue, we evaluated the effect of fixation on analyte detection, spatial localization, and introduction of artefactual signals. Hence, we assessed the molecular identity of lipids generated by matrix-assisted laser desorption ionization (MALDI-IMS) and liquid chromatography coupled tandem mass spectrometry (LC-MS/MS) for fixed and fresh-frozen retina tissues in positive and negative ion modes. Based on MALDI-IMS analysis, more lipid signals were observed in fixed compared to fresh-frozen retina. More potassium adducts were observed in fresh-frozen tissues than fixed as the fixation process caused displacement of potassium adducts to protonated and sodiated species in ion positive ion mode. LC-MS/MS analysis revealed an overall decrease in lipid signals due to fixation that reduced glycerophospholipids and glycerolipids and conserved most sphingolipids and cholesteryl esters. The high quality and reproducible information from untargeted lipidomics analysis of fixed retina informs on all major lipid classes, similar to fresh-frozen retina, and serves as a steppingstone towards understanding of lipid alterations in retinal diseases.

## 1. INTRODUCTION

MALDI-IMS is an analytical tool capable of mapping the distribution of lipids at the single cell level with high spatial accuracy and chemical specificity.^1^ In a typical workflow, MALDI-IMS detects exact masses of unknown molecules.^1–4^ In contrast, LC-MS/MS facilitates structural elucidation by generating molecularly-specific fragmentation spectra.^1–3,5,6^ Combining analyte localization with accurate mass measurements from MALDI-IMS^2,7,8^ with structural characterization by LC-MS/MS^7,9^ greatly enhances the breadth and depth of analysis of a tissue lipidome.^10–12^ MALDI-IMS analysis is typically done directly on fresh-frozen tissue sections so as to avoid chemical alterations.^13-21^ However, freezing can have deleterious effects on fine tissue morphology^13^. A few published studies investigated the effect of tissue fixation on lipid IMS^22–28^.

Paraformaldehyde-fixed paraffin-embedded fixation is the gold standard method for tissue preservation and histological examination.^29–31^ For MALDI-IMS analysis, fixed tissue sections are typically placed on poly-lysine coated indium-tin-oxide slides and deparaffinized with organic solvents, such as xylene, followed by matrix application^32^. A major limitation of fixed sections for MALDI-IMS is loss of sample lipids due to organic solvents not only at the deparaffinization step but also at the initial embedding steps. ^32–34^ An alternative approach is to fix tissue without embedding in paraffin, followed by cryosectioning^1^. However, fixation introduces chemical modification by cross-linking formaldehyde to amine/thiols in tissue biomolecules.^31^ Thus, it remains to be determined if the same lipids are detected in fixed and fresh-frozen tissues.

The retina is part of the central nervous system that separates from the brain in development and has its own neural circuits and unique lipids. ^35–37^ The lipid composition of human retina and the role of polyunsaturated long chain fatty acids and lipoprotein-related lipids of intraocular and systemic origin in age related macular degeneration (AMD) and other diseases is under active investigation. ^35,36,38,39 40,41^ The retina offers both opportunities and challenges for MALDI-IMS. The retina has exquisite tissue architecture due to horizontally alligned and structurally distinct layers of cells that are visible in histology and in diagnostic clinical imaging.^42^ However, the cone and rod photoreceptors are particularly delicate and prone to detaching from their underlying support tissues post-mortem. Furthermore, the fovea, a tissue specialization for high acuity and color vision, is small (<1 mm), thin (~250 μm), and easily damaged. Hence, to investigate this complex tissue at the molecular level with spatial accuracy, fixation is preferred to retain morphology.^23,43^

Previous studies demonstrated that lipids in fixed tissue differed from those in fresh-frozen tissue based on UPLC-MS, GC-MS, and MALDI-IMS technologies. ^22,32,34,44–50^ Of these three technologies, MALDI-IMS exhibited the maximal overlap of ions observed in paraformaldehyde-fixed and fresh-frozen tissue.^22–27^ Regarding studies of retina, Ly et al^23^ developed and compared lipid imaging by applying MALDI-IMS in fresh-frozen vs fixed mammalian (porcine) retina. To date molecular identification and characterization of lipids in fresh-frozen compared to fixed human retina has not been reported using MALDI-IMS. Further, extraction conditions and separation efficiency of LC-MS/MS to maximize lipidome coverage have not been explored for human retina. Herein, we performed an in-depth investigation of human retina and its supporting tissue using two complementary analytical methods, MALDI-IMS and LC-MS/MS in fresh-frozen and fixed human retina in positive and negative ion modes to determine if fixation adversely affects analyte detection or introduces artifactual signals.

## MATERIALS AND METHODS

### 2.1 Materials

HPLC grade chloroform, methanol, isopropanol, acetonitrile and water were purchased from Fisher Scientific (Pittsburg, PA). Formic acid and ammonium formate were obtained from Sigma-Aldrich (St. Louis, MO). tert-Butyl methyl ether (MTBE) was purchased from Honeywell (Charlotte, NC). SPLASH® LIPIDOMIX® Mass Spec Standard (15:0-18:1(d7) PC, 15:0-18:1(d7), PE, 15:0-18:1(d7) PS (Na Salt), 15:0-18:1(d7) PG (Na Salt), 15:0-18:1(d7) PI (NH4 Salt), 15:0-18:1(d7) PA (Na Salt), 18:1(d7) Lyso PC, 18:1(d7) Lyso PE, 18:1(d7) Chol Ester, 18:1(d7) MAG, 15:0-18:1(d7) DAG, 15:0-18:1(d7)-15:0 TAG, d18:1-18:1(d9) SM, Cholesterol (d7)), CL 18: 2(d5) and C24 Cer(d18:1/24:0) were from Avanti Polar Lipids, Alabaster, AL, U.S.

### 2.2 Methods

#### 2.2.1 Tissue preparation and MALDI-IMS procedure

Whole eyes were obtained from deceased human donors via Advancing Sight Network (Birmingham AL USA). As part of a study of AMD, donors were ≥ 80 years of age, white, and non-diabetic, and death-to-processing time was ≤6 hours. Tissues were prepared at the University of Alabama Birmingham and data acquisition was performed at Vanderbilt University. Methods were performed as described in Anderson *et al*. ^4^ In brief, the cornea was removed and the iris incised radially to allow fixative to infiltrate the globe. Globes were immersed in phosphate buffered 4% paraformaldehyde (PFA) for 24 hours, then in 1% PFA until processed. Fresh-frozen eyes were prepared without fixation and the cornea removed. The posterior part of the eye (sclera, choroid, retina) was dissected and embedded in carboxymethylcellulose (CMC) alongside the fixed eye so that the fresh-frozen and fixed eyes were in the same block, sectioned together, and sections placed on the same slide and processed for IMS together. CMC powder was mixed in deionized water, heated to 70°C, stirred until dissolved.^51^ Full-eye-wall thickness rectangular pieces of both fixed and fresh-frozen tissues were placed into cryomolds (Peel-A-Way 22 x 30 mm molds, Polysciences # 18646B) using a stereo microscope and oriented so the superior edge was sectioned first. Molds were filled with 15 ml of carboxymethyl cellulose, flash frozen with liquid nitrogen, and stored at −80°C until sectioned. For histopathologic evaluation, 12 μm cryosections were collected from both fresh-frozen and fixed tissues in the same tissue block. Sections were placed on pre-labeled 25 x 75 mm glass slides coated with a dilution of 0.01% poly-L-Lysine (Sigma Aldrich, St. Louis, MO, USA) and maintained at −20° during sectioning. During this process, 10 sections at 20 μm thickness for LC-MS were collected and thaw-mounted and dried onto precleaned uncoated glass slides (Tanner Scientific, FL, USA). At pre-defined intervals in the glass slide series, sections for IMS analysis were captured on large, 45×45 mm indium-tin-oxide slides (Delta Technologies Loveland, CO, USA), coated in-house with poly-lysine. A slide box was vacuum-packed with an oxygen-absorbing packet to help prevent oxidation and deterioration of lipid signal and stored at −80°C in a Bitran freezer bag for transport to Vanderbilt.

Samples were brought to room temperature before the matrices, 2,5-dihydroxyacetophenone (DHA) for positive ion mode analysis and 1,5-diaminonaphalene (DAN) for negative ion mode analysis, were applied using an in-house sublimation device set at 130 ° C. Sublimation times were 8 minutes for DHA and 15 minutes for DAN. MALDI-IMS data were acquired in positive and negative ion mode with a 15-20 μm pixel size in full scan using a solariX 9.4T FTICR mass spectrometer (Bruker Daltonics) equipped with a modified source to include an external desorption laser and associated optics to provide higher spatial resolution allowing for a 10 μm pixel size.

#### 2.2.2 MALDI-IMS data processing

MALDI-IMS data were processed using the R package *Cardinal*^52^. Data were loaded from the peak picked Bruker .sqlite file with a mass resolution of ppm = 0.5. Using *Cardinal*, a mean spectrum was generated from across all spectra and peak picking (signal-to-noise ratio = 3) was performed to reduce the data to 618 and 329 features in positive and negative modes, respectively. Exact masses of these features were searched in LIPIDMAPS, and tentative annotations were made from database hits. Tentative annotations were then collated by lipid class for comparison across preparation methods. Differential analysis of frozen and fixed data was performed by comparing mean spectra after removal of off-tissue pixels. Noise and off-tissue IMS pixels were identified by using the *spatial shrunken centroids* algorithm in *Cardinal* (using parameters k = 10, r=1 and s=1)^53^ and removing segments with primarily matrix signal in the *t-statistics* plot.

#### 2.2.3 Lipid extraction and LC-MS/MS procedure

Three human donor eyes identified and processed as described above were used for lipid extraction. Ten samples of 20 μm thick sections of embedded fresh-frozen and fixed retina tissue were used for lipid extraction. Under a dissecting microscope, the sclera was carefully removed with a scalpel blade. The remaining retina-choroid tissue was scraped from the glass slide and placed in a 1.5 mL HPLC glass vial. One mL of MMC^54^ extraction mixture (1.3:1:1, methanol: tert-Butyl methyl ether (MTBE): chloroform) was spiked with 10 μL of lipid standard mixture (SPLASH® LIPIDOMIX® Mass Spec Standard from Avanti Polar Lipids, Alabaster, AL, U.S) in the glass vial, vortexed for 60 seconds and centrifuged at 3000 rpm for 10 minutes. The supernatant was transferred to a separate HPLC vial and was evaporated to dryness. The sample was then reconstituted with 30 μL methanol and 10 μL was subjected to LC-MS/MS analysis.

Chromatographic separation^5^ was achieved using an ACQUITY UPLC BEH C-18 column (2.1 X 150mm, 1.7 μm particle size) (Waters Corporation, Milford, MA, USA) held at 50°C coupled to a Vanquish binary pump (Thermo Fisher Scientific, San Jose, CA). A gradient mobile phase was generated over 55 minutes with a flow rate of 250μL/min and was comprised of 10 mM ammonium formate in 40:60 (v/v) water: acetonitrile with 0.1% formic acid and 10 mM ammonium formate in 90:10 v/v) isopropanol: acetonitrile with 0.1% formic acid. The gradient elution profile was as follows: 32% B (0-1.5 min), 32-45% B(1.5-4 min), 45-52% B(4-6 min), 52-58% B(6-10 min), 58-66% B (10-16 min), 66-70% B (16-22 min), 70-75% B(22-30 min), 75-97% B(30-38 min), 97%B (38-48 min), 97-32% B(48-50 min), 32% B(50-55 min). Ten μL of sample was injected via the Vanquish autosampler maintained at 4°C, HPLC eluate flowed into a HESI heated ESI source, and ions were analyzed using a Q Exactive HF instrument (Thermo Scientific, San Jose, CA, USA). Data were acquired at both full (MS1) and data dependent MS2 (ddMS2) scan modes using positive and negative modes separately. The full scan mode had a mass resolution of 60000, mass range of m/z 400-1200 in positive polarity and m/z 240-1600 in negative polarity and a maximum trap fill time of 100 ms. ddMS2 data were acquired at 15,000 resolution with a maximum trap fill time of 160ms. The isolation window of selected MS1 ions was ± 1.4 m/z with a normalized collision energy (NCE) of 20 and 25. LC-MS/MS data were acquired using Xcalibur version 4.0.

#### 2.2.4 Data analysis

Raw LC-MS/MS data files (Thermo .raw files) were converted to mgf, mzXML files using the MSConvert function of ProteoWizard. Mgf, mzXML input files were searched using the spectrum searcher feature of LipiDex software v1.0.2^55^ for lipid identifications using LipiDex_HCD_Formic and LipiDex_Splash_ISTD_Formic libraries. Precursor mass-to-charge (m/z), fragmentation spectra, chromatographic retention time and identification were integrated using LipiDex. Fragmentation patterns of individual lipid standards were manually interpreted and then matched against LipiDex library results.

To ensure correct annotation of individual molecular ion species, accurate m/z values for the precursor ions and subsequent ddMS2 spectra were acquired from the SPLASH® LIPIDOMIX® mixture of lipids. Lipid species were identified based on their fragmentation pattern with a mass tolerance of 0.2 ppm in negative and positive ionization modes. Confirmed lipid identifications were made by examination of tandem mass spectra for head group and fatty acyl chains, mass error measurement, isotopic profile, extracted ion chromatogram, and the retention time for precursor ions. LC-MS/MS data were processed to separate each lipid identification into its distinct lipid class. For each replicate, the sum of unique IDs in each lipid class was determined and set unions were calculated in custom R scripts. For m/z vs retention time (rt) plotting, the earliest rt signal of each unique ID was used for visualization.

## RESULTS AND DISCUSSION

### 3.1 Comparison of fresh-frozen and fixed human retina lipid detection by MALDI-IMS

In MALDI-IMS (**Figure 1)**, a total of 358 lipid signals were identified including 299 in fresh-frozen and 329 in fixed retina based on MS1 confirmation including negative and positive ionization modes. There was a 10% higher number of lipid signals observed for fixed tissue as compared to fresh-frozen. There are potentially more signals observed in the fixed tissue as lower abundance molecular signals are no longer spilt between protonated, sodiated and potassiated species resulting in higher signal to noise, allowing them to be selected in the peak picking process. Other possible explanations for this observation are that fixation helps in preservation of tissue morphology limiting ion suppression and that less molecular degradation may occur in fixed tissue compared to fresh-frozen tissue^56,57^. Further studies are required to confirm these possible reasons for the observed increased number of lipid signals in fixed retina tissue.

**Figure 1:**
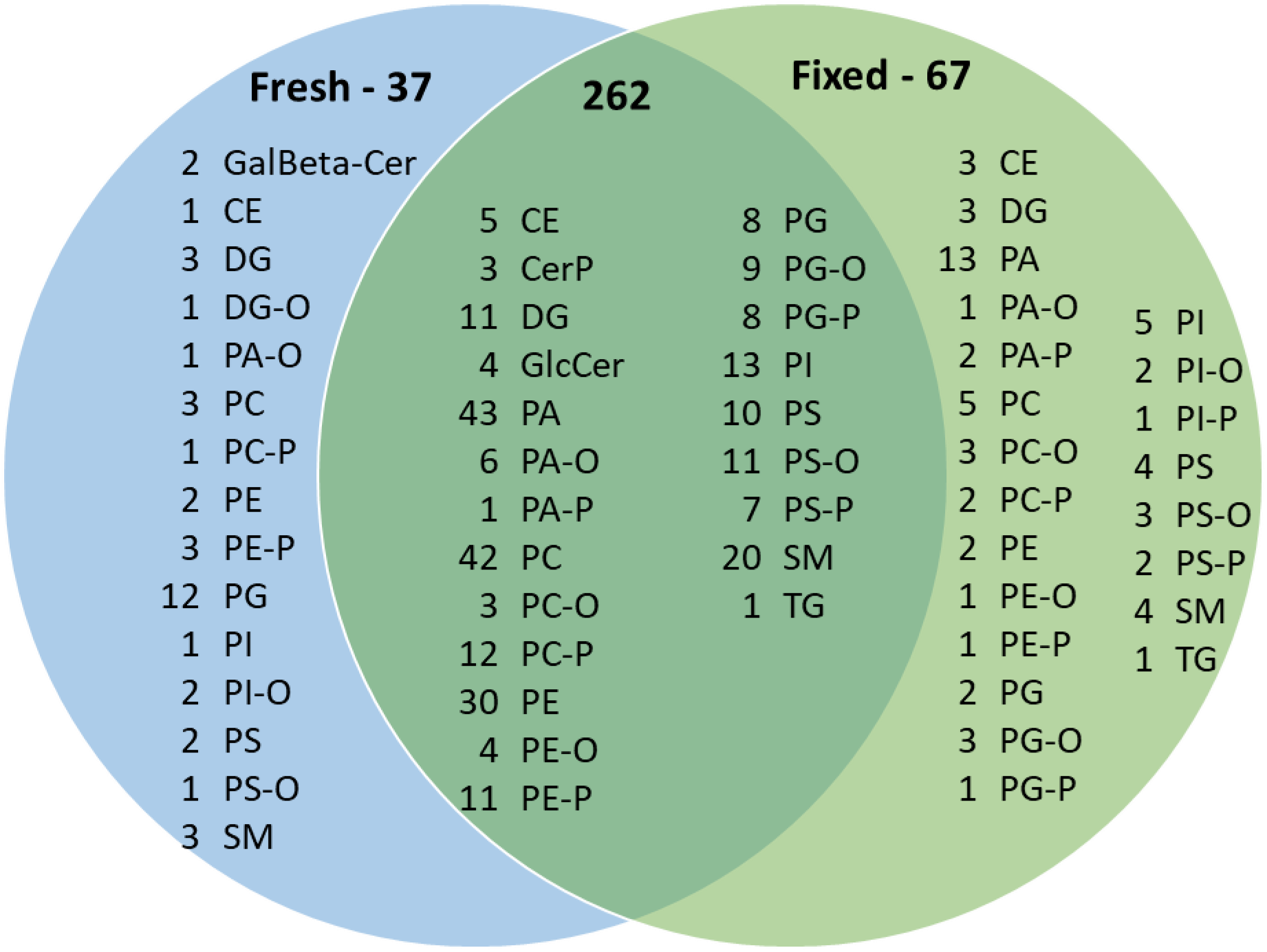
Venn diagram of lipid compositions of fresh-frozen and fixed human retina including both negative and positive ionization modes detected by MALDI-IMS.

Most lipid subclasses were conserved upon fixation including glycerolipids, sphingolipids and cholesteryl esters with the exception being glycerophospholipids (**Figure 1**). The major classes of lipids observed in IMS data include: glycerophospholipids; PC, PE, PG, PI, PS, PA, PC-O, PC-P, PE-O, PE-P, PG-O, PG-P, PS-P, PA-O, PA-P, PS-O, PS-P (217 common, 26 in fresh-frozen only and 51 in fixed only), glycerolipids; DG, DG-O, DGTS, MGDG, TG (14 in common, 2 in fresh-frozen only and 2 in fixed only), sphingolipids; SM, GalBeta-Cer, CerP, (32 common, 5 in fresh-frozen only and 3 in fixed only), cholesteryl esters (4 common, 1 in fresh-frozen only and 4 in fixed only). The variations in glycerophospholipid signals were mainly observed for PG/BMP species. More PA lipids were detected in fixed tissues

### 3.2 Lipid signal intensities for fresh-frozen and fixed retina in positive and negative ionization modes using MALDI-IMS

In this section and in Figure 2 and Supplementary Figures 1-4, “retina” includes “inner retina” and “outer retina”. Inner retina includes nerve fiber layer, ganglion cell layer, inner plexiform layer, and inner nuclear layer (NFL, GCL, IPL, INL). Outer retina includes outer plexiform layer, outer nuclear layer, inner and outer segments of photoreceptors (bacillary layer), and retinal pigment epithelium (OPL, ONL, IS and OS, RPE). Choroid is a vascular bed behind the retina, and sclera is behind the choroid.

**Figure 2:**
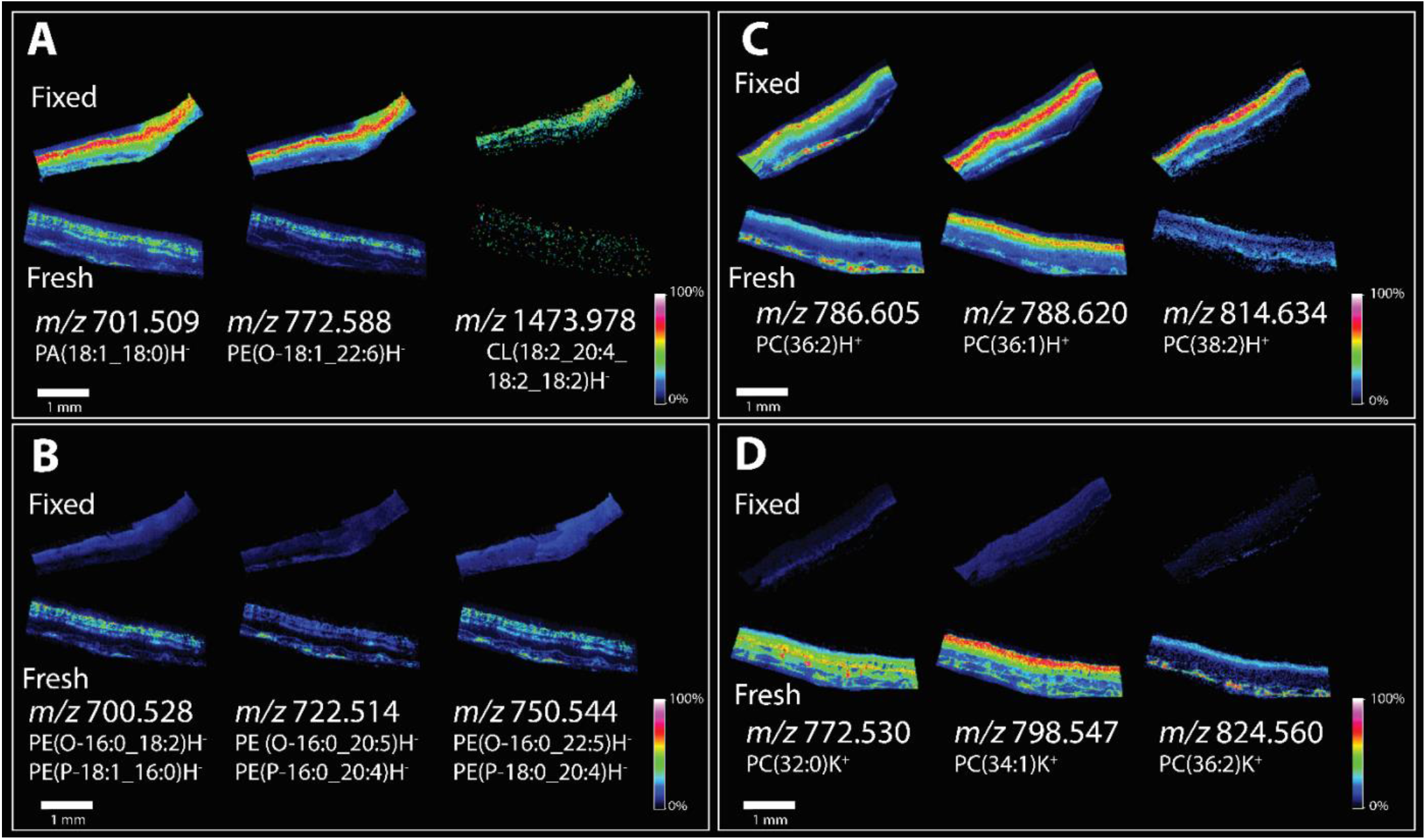
MALDI-IMS analysis of 90-year-old human donor eye from fresh-frozen sample vs paraformaldehyde-fixed in peripheral sections from the same donor. Panels A-D display multiple lipid signals in both polarities in fixed and fresh-frozen tissue. Panel A and B displays negative ion mode analysis were three signals which were observed to have higher signal intensity in fixed tissue (A) and fresh-frozen tissue (B). Panel C and D displays positive ion mode analysis were three signals which were observed to have higher signal intensity in in fixed tissue (C) and fresh-frozen tissue (D).

Lipid signal variations in IMS analysis of fresh-frozen and fixed tissue are demonstrated in **Figure 2**. The negative ion data show the highest intensity for most lipid signals in fixed tissue. Three examples of these lipid signals are shown in **Figure 2A**. The investigated lipids PA, PE and CL were observed with similar distributions in the inner retina of the fixed tissue. **Supplementary Figures 1 and 2** display the same signals from **Figure 2** overlaid with optical images to clearly indicate signal localization in the tissue and reproducibility of the results. Three examples of lipid signals that were slightly higher in negative ion mode for fresh-frozen tissue than for fixed tissue are shown in **Figure 2B**. These signals were comprised of plasmalogen and ether-linked PE lipids that have the same mass and cannot be distinguished from accurate mass alone; both were detected by LC-MS/MS. These signals were observed both in the retina and choroid.

In positive ionization mode the tentative identifications suggest a higher abundance of protonated species in fixed tissues as compared to fresh-frozen tissue, localized to the inner and outer retina for PC(36:2)H^+^ and PC(36:1)H^+^ and the inner retina for PC(38:2)H^+^ (**Figure 2C**). Variations in signal intensity for the positive ion mode analysis can be partially attributed to adduct formation in fresh-frozen tissue **(Figure 2D)**. High abundance of potassium adducts can be seen in the fresh-frozen inner retina for PC(32:0)K^+^, PC(34:1)K^+^ and in the choroid, PC(36:2)K^+^. Very low signals for these adducts were observed in fixed tissue. A shift from potassium to sodium adduct formation has been previously observed by Carter *et al*^24^ in fixed brain tissue. The distributions of protonated, sodiated and potassiated lipids shown in **Supplementary Figure 3** are all similar but vary in intensity.

Replicate MALDI-IMS data in **Supplementary Figure 4** clearly demonstrate the distorted morphology of fresh-frozen retina, underscoring the value of chemical fixation for this delicate tissue. Variations in CL signals were observed between biological replicates in **Figure 2** and in **Supplementary Figure 1**. The CL with high intensity in the fixed tissue in the first replicate **(Figure 2)** did not generate a quality IMS image in second replicate **(Supplementary Figure 1)**, although a peak was present. A similar observation was also made with PC(38:2) at *m/z* 814.634. An increase in protonated signal was less apparent in these data yet a trend toward high potassium adducts in fresh-frozen tissues was observed.

MALDI-IMS-investigated lipid signals based on accurate mass measurement in positive and negative ionization modes for fresh-frozen and fixed tissue are shown in **Supplementary Figure 5**. The protonated, sodiated and potassiated signals from individual lipid were counted as one signal.

Higher signal intensities in fixed tissues for most species could be a result of the improved morphology and preservation of multi-layered retina tissue. Fixed tissue may have reduced analyte suppression as compared to fresh-frozen in discrete locations as the layers and the native dimensions are preserved rather than disrupted and merged.

### 3.3 Elution profile of lipid classes extracted using LC-MS/MS

To compare LC-MS/MS lipidomic elution profiles we aligned the common lipids found in fresh-frozen and fixed tissue based on retention time vs precursor mass as shown in **Figure 3** from one of three biological replicates. **Supplementary Figures 7 and 8** show consistent results from two additional biological replicates.

**Figure 3:**
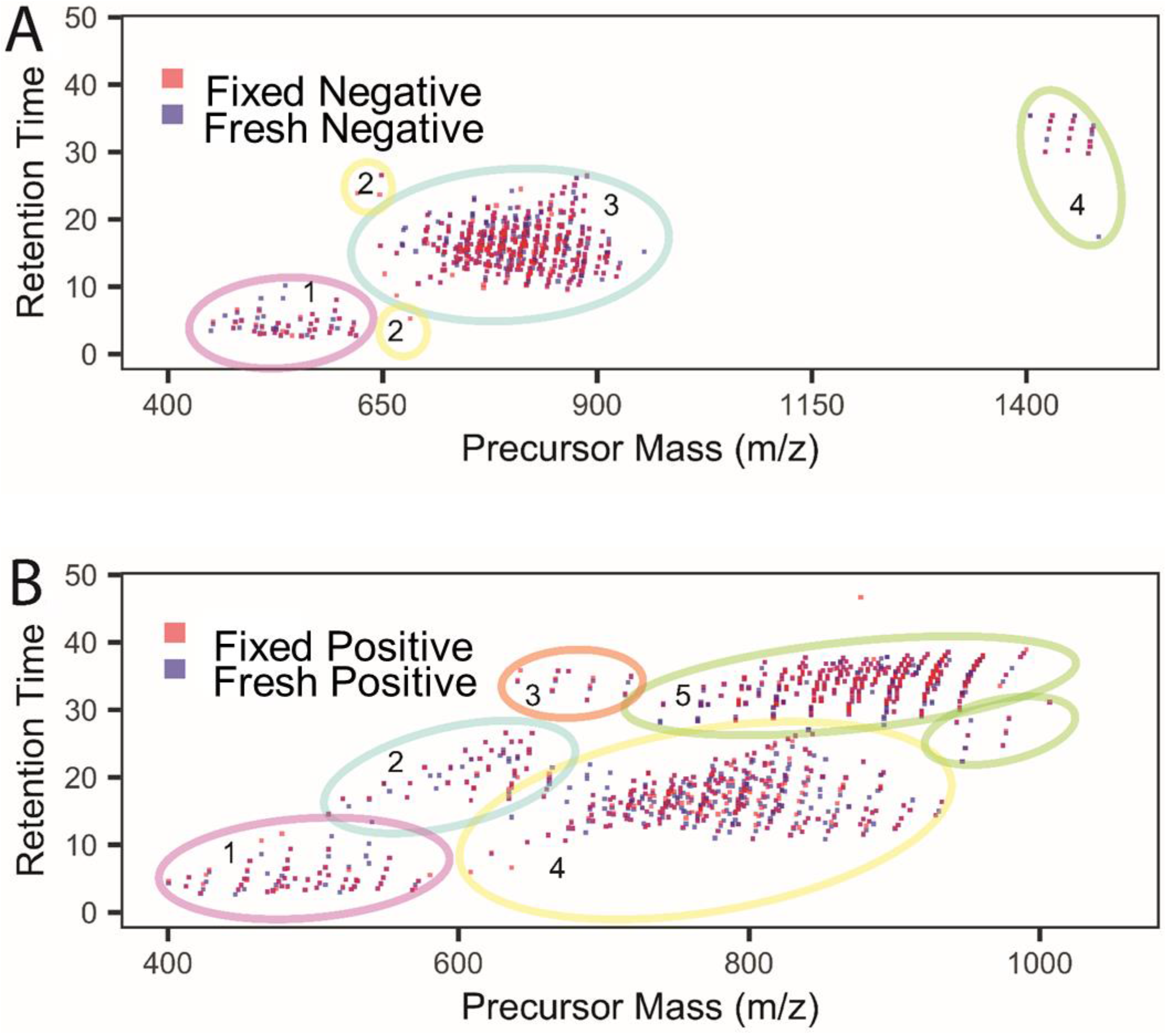
Summary of LC-MS/MS data of fresh-frozen (blue boxes) vs fixed (red boxes) human donor retinas. **(A)** m/z versus retention time summary of negative ion mode data where lipid classes were segregated (colored circles 1-4) based on their elution profiles. 1; lysoPC, lysoPE, lysoPG, lysoPI, lyso PS, 2; Cer, 3; PC,PE,PG,PS,PI,PA,PC-O, PE-O,SM, 4; CL. **(B)** M/z versus retention time summary of positive ion mode data where lipid classes were segregated (colored circles 1-5) based on their elution profiles. 1; AC, lyso PC, lyso PE, 2; Cer, PG, 3; CE, 4; PC, PE, PC-O, PC-P, PE-O, PE-P, PE-NMe; 5; TG.

In negative ionization mode lipids were stratified by retention time vs precursor mass into four groups (**Figure 3A**). In group 1, we observed lipids in the range of m/z 450-620 with retention times of 2-10 minutes and comprised of lyso PC, lyso PE, lyso PG, lyso PI and lyso PS. There were some outliers, i.e., lipids that did not segregate with groups 1 or 3, comprised of 4 distinct ceramides out of which 3 were exclusively present in fixed retina and 1 was common for both fresh-frozen and fixed tissue. These lipids were placed in group 2. Lipids eluting between 10-30 minutes within a m/z range of 650-900 were placed in group 3 and consisted mostly of glycerophospholipids including PC, PE, PG, PS, PI, PA, plasmenyl-PC, plasmenyl-PE and sphingomyelins (SM). Cardiolipins (CL) eluted last from the column, after 30 minutes, with masses above m/z 1400; they formed group 4.

In positive ionization mode (**Figure 3B**) lipids were sorted into 5 groups based on m/z and retention time. AC, lyso PC, lyso PE eluted first from the column at 2.6-9.4 minutes within a range of m/z 400-600 and were marked as group 1. Within the retention time range from 10-30 minutes eluted lipids in group 2, including Cer[NS] and DG within a mass window of m/z 500-700. Group 3 lipids, between m/z 640-728, included cholesteryl esters (CE) that eluted between 30-40 minutes. Glycerophospholipids including PC, PE, plasmenyl-PC, plasmenyl-PE, plasmanyl-PC, PE-NMe and sphingomyelins have m/z values between 650-950 were combined in group 4 with elution times of 10-30 minutes. Group 5, comprised of triglycerides (TG), eluted from 28.1-38.4 minutes with m/z values between 738-990. Interestingly, a discrete pattern was observed for TG lipids showing an increase in retention time with a proportionate increase in the degree of unsaturation.

### 3.4 Comparison of lipid composition of fresh-frozen and fixed human retina-choroid in negative and positive ionization modes using LC-ESI-MS/MS

Overall, untargeted lipidomic analysis in positive and negative ionization modes from three biological replicates identified a total of 592 lipids covering 4 lipid classes and 17 lipid subclasses comprised of 568 lipids in fresh-frozen retina (201 fresh-frozen only) and 391 lipids in fixed retina (24 fixed only) representing a 31% loss of unique lipids upon fixation (**Figure 4)**. Overall, there was 22%, 12% and 32% loss of lipids in fixed tissue as compared to fresh-frozen from three biological replicates, respectively. The majority of lipids discovered in both fresh-frozen and fixed tissues were glycerophospholipids, followed by glycerolipids, sphingolipids and cholesteryl esters. The highest number of detected lipid subclasses were diacylglycerophosphocholines (PC, 124), triacylglycerol (TG, 94), sphingomyelins (SM, 42), and cholesteryl esters (CE, 9). There were 352 glycerophospholipids in fresh-frozen and 250 in fixed tissue representing a 29% decrease, covering lipid subclasses including, PC, PI, PG, PE, PS, plasmalogen lipids and lyso phospholipids. A total of 114 glycerolipids, comprised of TGs and DG, were observed in fresh-frozen and 56 in fixed tissue; a 51% decrease. Sphingolipids were 94 in number for fresh-frozen and 79 for fixed tissue; a 16% decrease, including Cer, CL, SM, AC and HexCer. Upon fixation, lipid signals were preserved for most of the lipid classes. However lower signals were observed for PC, PE, PS, PI, PG, TG, AC and SM upon fixation (**Figure 4**).

**Figure 4:**
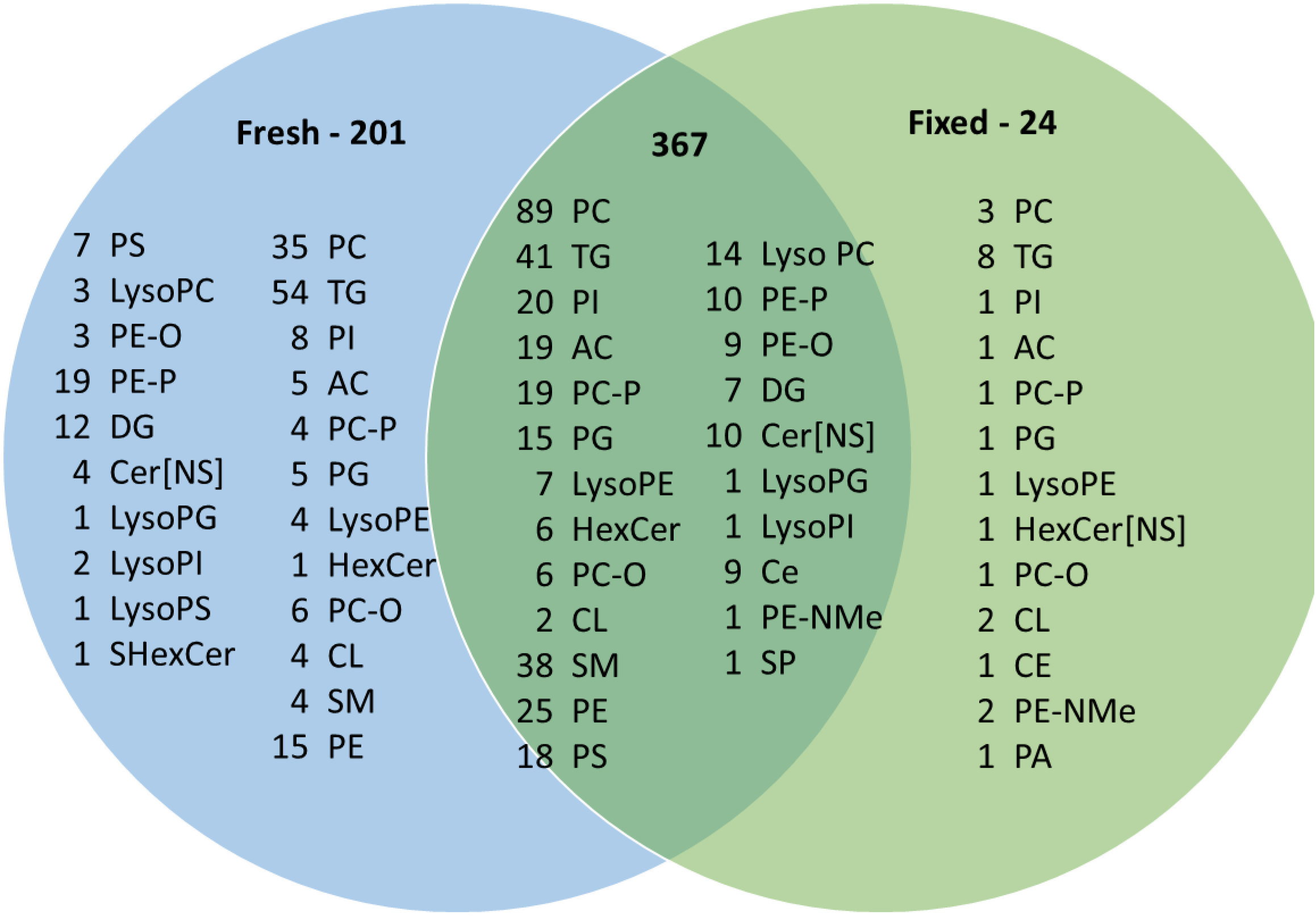
Venn diagram of lipid compositions of fresh-frozen and fixed human retina samples including both negative and positive ionization mode data identified by LC-MS/MS (only MS/MS confirmed lipids shown).

A comparison of overall differences in the lipid composition among three biological replicates of fresh-frozen and fixed tissue in negative ion mode **(Figure 5)** showed higher numbers of CL, PS, Lyso PLs, PE, PI and SM lipids were detected for fresh-frozen than for fixed samples. Fewer Cer, PA, PC, PG, plasmenyl PC lipids were detected in fresh-frozen as compared to fixed tissue. In positive ionization mode AC, Cer, DG, lyso PL, PC, PE, PI, plasmenyl PE, SM and TG lipids were detected in higher numbers from fresh-frozen tissue compared to fixed tissue and CE, SHexCer were detected in lower numbers in fresh-frozen compared to fixed tissue **(Figure 5**).

**Figure 5:**
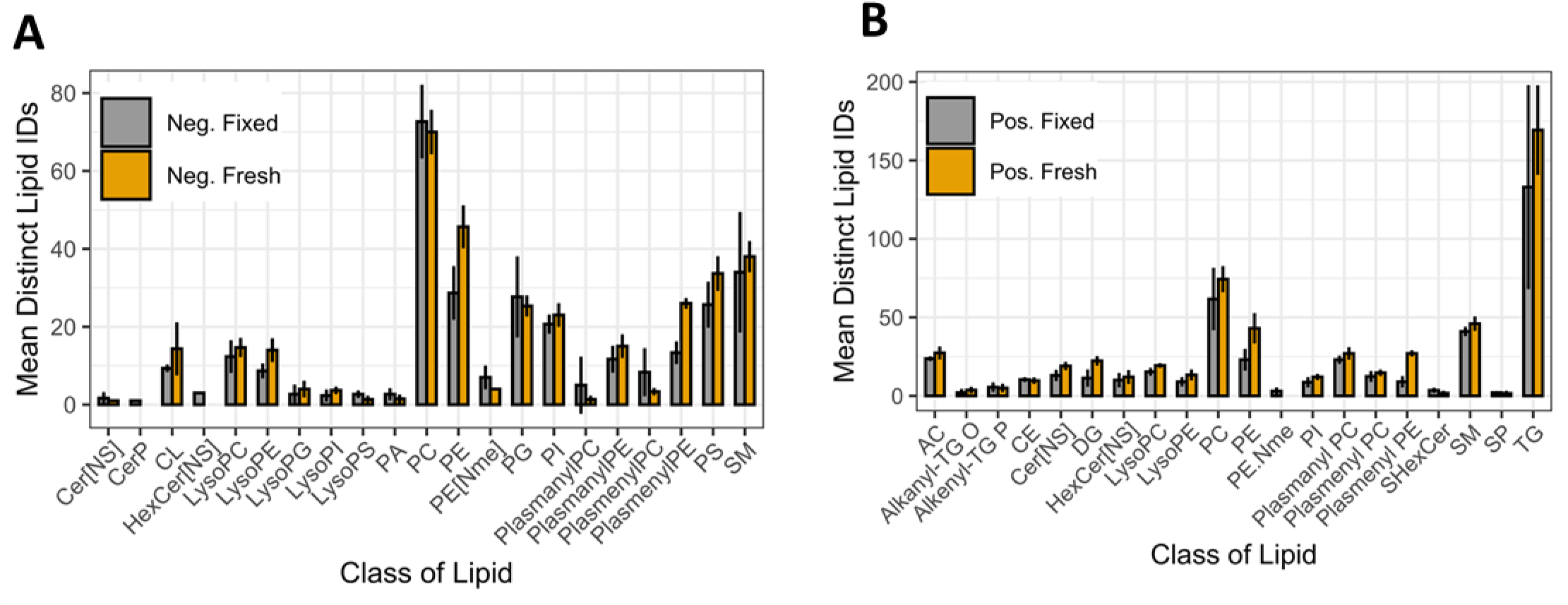
Differences in the lipid composition observed between fresh-frozen (orange) and fixed (gray) retina tissue samples detected by LC-MS/MS analysis in negative (A) and positive (B) ion modes (n=3).

Among glycerophospholipids, formalin reacts readily with the primary amine groups of PE and PS lipids to form a crosslink, thereby reducing the detection of free lipids in fixed tissue, as previously observed.^26,27,58^ Similarly, we observed a decrease in lipid signals for PE and PS lipids in all three biological replicates of fixed human retina **(Supplementary Figure 9)**. Also, monomethylation and dimethylation of the amine head group of PE by the nucleophilic addition of formaldehyde on primary amines leads to increased expression of PE-NMe in fixed samples (**Figure 4**). PC lipids were most affected by fixation and this can be attributed to lipid peroxidation^59^ and hydrolysis of PC^58^, as observed for PE,PS and PG (**Figure 4**). Fixation caused a decrease in lipid signals for PG lipids in our study. This contrasts with one study that showed no effect on PG lipids, ^27^ but supports two other studies.^26,58^ The slight increase in free fatty acid PA in the biological replicates of fixed retina could be attributed to hydrolysis of the ester bond linking the fatty acyl chain. PI lipids showed decreased signals due to fixation which is consistent with other reports^27^. The reduction in PI levels may be due to either minor degradation of endogenous lipids during the fixation process and/or partial reaction of the hydroxyl groups of PI with formalin^27^. Plasmalogen lipids originating from PC, PE, PI and PS had decrease signals in fixed tissue which is attributed to degradation of vinyl ether linkage at the sn-1 position of these species.

Glycerolipids, including TG and DG were also decreased due to fixation as shown in **Figure 4**. Comparing fresh-frozen and fixed tissue from biological triplicates, a large increase in number of lipids for TG and DG in fresh-frozen tissue was observed in one sample (**Supplementary Figure 9**), while the other two samples showed a slight decrease. A previous study using tissue of one representative human brain showed a decrease in DG lipids in fixed tissue that was attributed to chemical hydrolysis of phosphodiesterase bonds^58^.

Sphingolipid signals were mostly preserved after fixation (**Figure 4**). A comparison of three biological replicates indicated there were fewer CL observed in fixed tissue (**Supplementary Figure 9**). Among sterols, cholesteryl esters (CE) were observed and are very stable, unlikely to be affected by fixation. CE signals were conserved throughout the samples of fresh-frozen and fixed retina in all the three biological replicates. Previous studies found that longer chain CE lipids became less detectable in fixed tissue.^60,61^

## CONCLUSIONS

Fixation to preserve tissue morphology for MALDI-IMS analysis, in conjunction with LC-MS/MS to identify molecular species observed with MALDI-IMS, together can elucidate lipid localization, utilization and accumulation in healthy and diseased retina. Extensive untargeted lipidomics analysis using the analytical discovery tools of LC-MS/MS and MALDI-IMS revealed that molecular signatures for most lipid classes in human retina tissue were preserved after fixation. However, fixation causes lipid crosslinks, oxidation, hydrolysis, and head group modifications for certain lipids. There was a 31% overall decrease in lipid signals for fixed tissue based on LC-MS/MS analysis. Fixation, compared to fresh-frozen tissue results, caused decreased lipid signals for most of glycerophospholipids and glycerolipids, and increased lipid signals for PG, PE-NMe and plasmalogen PC lipids. Most sphingolipids and cholesteryl esters were unaltered as detected by LC-MS/MS. Based on MALDI-IMS, fixation caused an increase in signals for most of the lipid classes in negative ionization mode while an increase in signal was observed for PG/BMP and PS in positive ionization modes. Increased signals observed in fixed tissue is likely due to reduced ion suppression effects in tissues where morphology of retina layers is preserved. The morphology of the retina was observed to be distorted in fresh-frozen tissue sections and this suggests caution should be used for this sample type. Formation of abundant potassium adducts in fresh-frozen tissue and displacement of these adducts in fixed tissue increased the number of signals detected in fresh-frozen tissue but did not increase the number of unique lipids identified. Overall, our results suggest that fixed tissue, typically used to maintain tissue morphology, conserves lipid classes as seen in fresh-frozen tissue and can be used to investigate retina lipids in various retinal disease settings.

## ACKNOWLEDGEMENTS

The authors acknowledge financial support from NIH grants R01EY027948 (CAC), P41 GM103391, S10 OD023514 (KLS), Heidelberg Engineering, (UAB institutional support), Research to Prevent Blindness (CAC & KLS), and EyeSight Foundation of Alabama.

## Supplementary legends

**Supplementary Figure 1 and 2:** MALDI-IMS analysis of 90-year-old human donor eye from fresh-frozen sample vs paraformaldehyde-fixed in peripheral sections from the same donor. Panels A-D display multiple lipid signals in both polarities in fixed and fresh-frozen tissue overlaid with an optical image of the tissues imaged. Varying signal intensities from the selected species can be seen for both fresh-frozen and fixed tissue in both polarities, Supplement Figure 1 for negative ion and Supplement Figure 2 for positive ion.

**Supplementary Figure 3:** Positive ion mode MALDI-IMS of fresh-frozen vs fixed tissue displaying **A** PC(32:0), **B** PC(34:1) and **C** PC(34:2) from Figure 2D displaying the protonated, sodiated and potassiated ions which show the same localization but varying intensities.

**Supplement Figure 4:** Replicate **i**maging mass spectrometry (IMS) data analysis of 84 year old human donor eye from fresh-frozen sample vs paraformaldehyde-fixed in peripheral sections from the same donor. Panels A-D display multiple lipid signals in both polarities in fixed and fresh-frozen tissue. Panel A and B display negative ion mode analysis were three signals which were observed to have higher signal intensity in in fixed tissue (A) and fresh-frozen tissue (B). Panel C and D display negative ion mode analysis were three signals which were observed to have higher signal intensity in in fixed tissue (C) and fresh-frozen tissue (D). The poor morphology of the fresh-frozen tissue if reflected in the image quality in this example.

**Supplement Figure 5:** Panel A and B show the differences in lipid composition from fixed (gray bars) and fresh-frozen (yellow bars) tissues analyzed using MALDI-IMS displaying multiple lipid classes observed in negative (Panel A) and positive (Panel B) ion mode analysis.

**Supplementary Figure 6 A**: Molecular ion spectra and fragmentation pattern in negative ionization mode of fixed tissue using LC-ESI-MS/MS. Calculated monoisotopic mass **A1**. 701.5121; PA (18:1_18:0); ppm error: 1.0, **A2**. 772.5281; PE(O-18:1_22:6); ppm error: 0.6 and **A3**. 1473.981; CL (18:2_20:4_18:2_18:2); ppm error: 1.3. **B**: Molecular ion spectra and fragmentation pattern in negative ionization mode of fresh-frozen tissue using LC-ESI-MS/MS. Calculated monoisotopic mass **B1**. 700.5281; **(a)** PE(O-16:0_18:2) and **(b)** PE(P-18:1_16:0); ppm error: 0.1, **B2**. 722.5125; **(a)** PE(O-16:0_20:5) and **(b)** PE(P-16:0_20:4); ppm error: 0.8, **B3**. 750.5443; **(a)** PE(O-16:0_22:5) and **(b)** PE(P-18:0_20:4); ppm error: 0.3. **C:** Molecular ion spectra and fragmentation pattern in positive ionization mode of fixed tissue using LC-ESI-MS/MS. Calculated monoisotopic mass **C1**. [M+H]+ 786.6013; PC(36:2); ppm error:2.6, **C2**. [M+H]+ 788.6169; PC(36:1); ppm error:0.6, **C3**. [M+H]+ 814.6326; PC(38:2); ppm error: 5.6. **D**; Molecular ion spectra and fragmentation pattern in positive ionization mode of fixed tissue using LC-ESI-MS/MS. Calculated monoisotopic mass **D1**. [M+H]+ 734.5694; PC(32:0); ppm error: 1.9, **D2**. [M+H]+ 760.5851; PC(34:1); ppm error: 2.2, **D3**. [M+H]+ 786.6007; PC(36:2); ppm error: 1.9.

**Supplementary Figure 7:** LC-MS/MS data of fresh-frozen (blue boxes) vs fixed (red boxes) human donor retina. **(A; biological replicate 2 and B; biological replicate 3)** m/z versus retention time summary of **(a)** negative ion mode data and m/z versus retention time summary of **(b)** positive ion mode data where lipid classes.

**Supplementary Figure 8:** Lipid composition identified (MS2 confirmations) in fresh-frozen and fixed human retina from LC-MS/MS using LipiDex.

