## Supplemental Figures and Table for "Tissue fixation effects on human retinal lipid analysis by MALDI imaging and LC-MS/MS technologies"

Ankita Kotnala[1,2], David M.G. Anderson[1], Nathan Heath Patterson[1], Lee S. Cantrell[1], Jeffrey D. Messinger[2], Christine A. Curcio[2] and Kevin L. Schey[1].

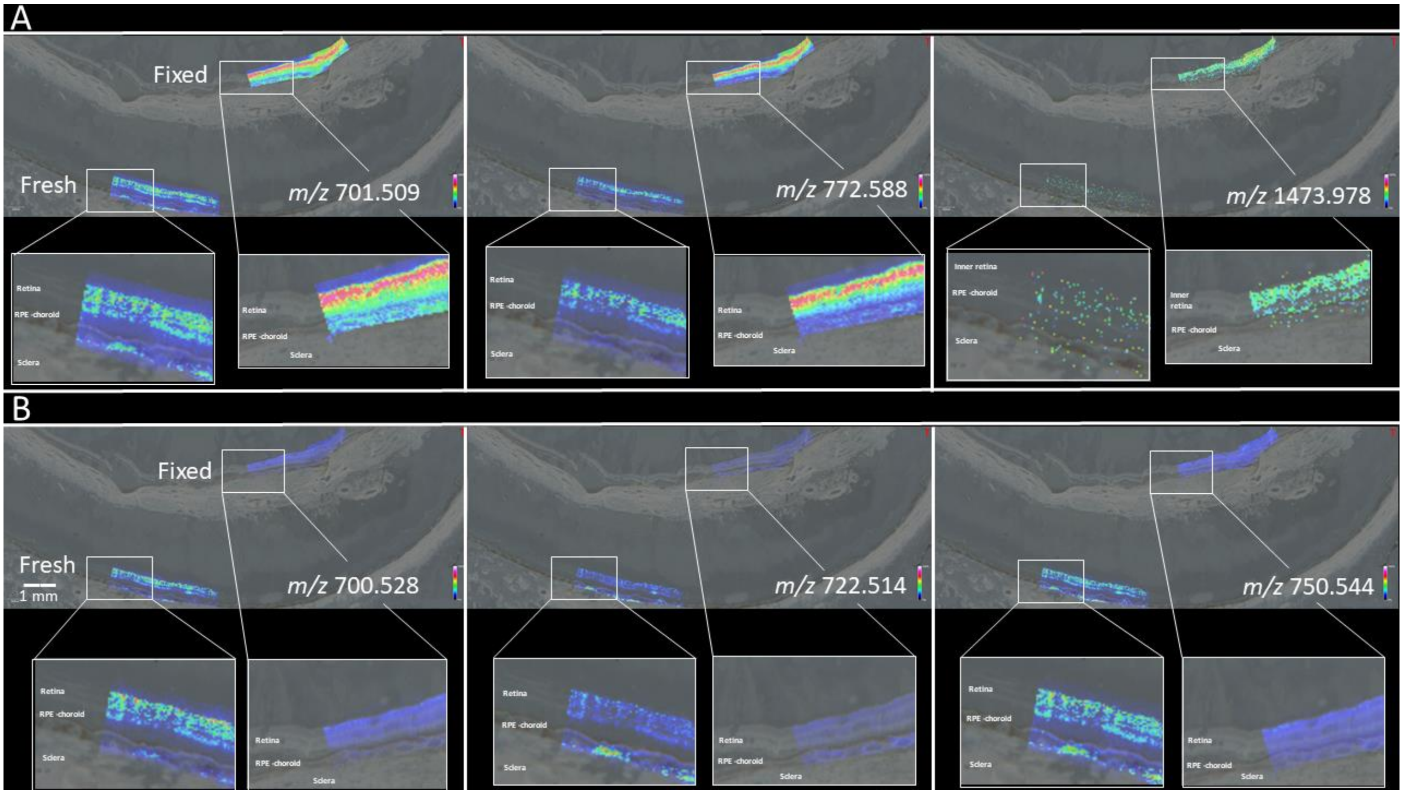


**Supplementary Figure 1:** Negative ion MALDI-IMS analysis of 90-year-old human donor eye from fresh-frozen sample vs paraformaldehyde-fixed in peripheral sections from the same donor. Panels display multiple lipid signals in in fixed and fresh-frozen tissue overlaid with an optical image of the tissues imaged. Varying signal intensities from the selected species can be seen for both fresh-frozen and fixed tissue.

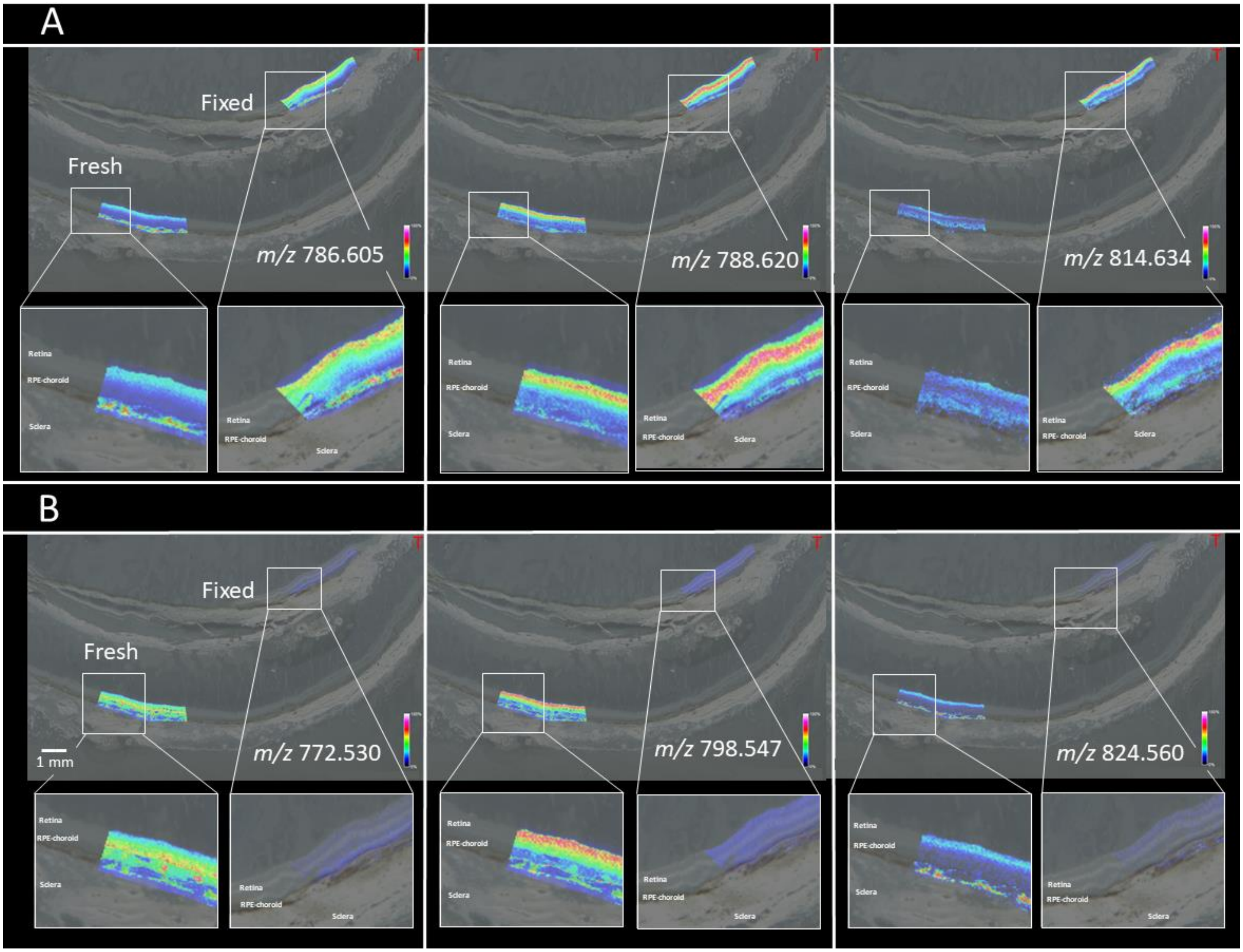


**Supplementary Figure 2:** Positive ion MALDI-IMS analysis of 90-year-old human donor eye from fresh-frozen sample vs paraformaldehyde-fixed in peripheral sections from the same donor. Panels display multiple lipid signals in in fixed and fresh-frozen tissue overlaid with an optical image of the tissues imaged. Varying signal intensities from the selected species can be seen for both fresh-frozen and fixed tissue.

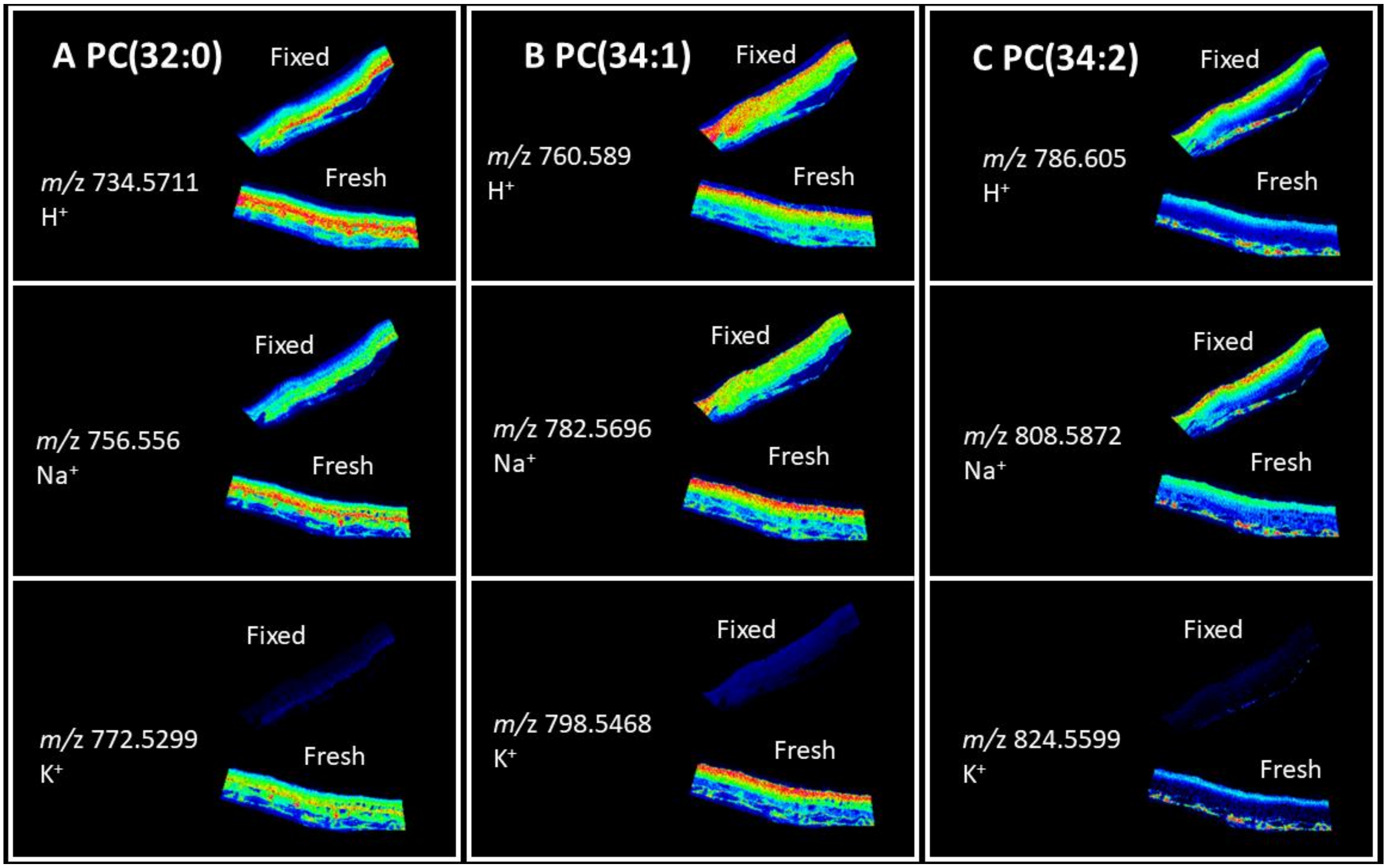


**Supplementary Figure 3:** Positive ion mode MALDI-IMS of fresh-frozen vs fixed tissue displaying **A** PC(32:0), **B** PC(34:1) and **C** PC(34:2) from Figure 2D displaying the protonated, sodiated and potassiated ions which show the same localization but varying intensities.

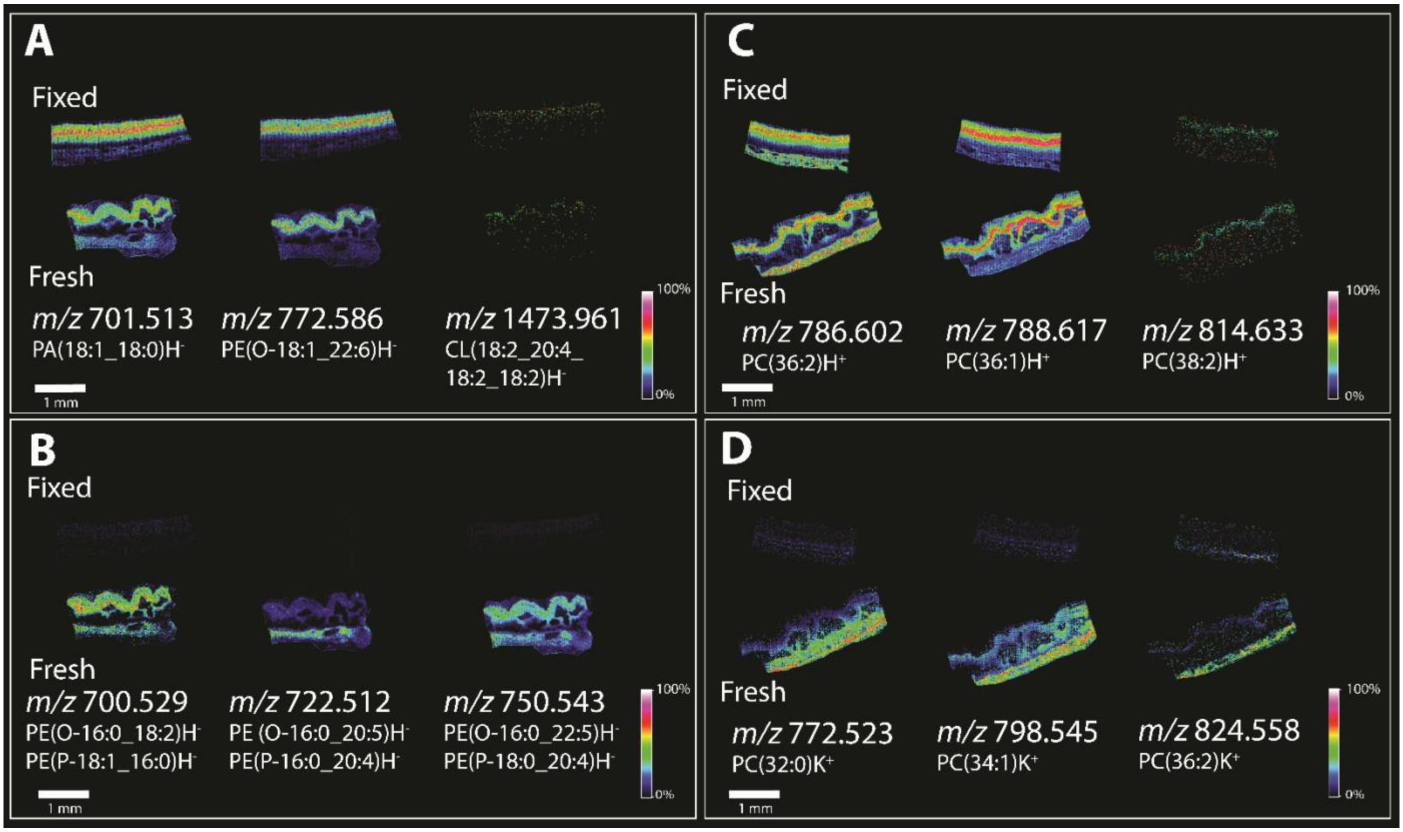


**Supplement Figure 4:** Replicate imaging mass spectrometry (IMS) data analysis of 84 year old human donor eye from fresh-frozen sample vs paraformaldehyde-fixed in peripheral sections from the same donor. Panels A-D display multiple lipid signals in both polarities in fixed and fresh-frozen tissue. Panel A and B display negative ion mode analysis were three signals which were observed to have higher signal intensity in in fixed tissue (A) and fresh-frozen tissue (B). Panel C and D display negative ion mode analysis were three signals which were observed to have higher signal intensity in in fixed tissue (C) and fresh-frozen tissue (D). The poor morphology of the fresh-frozen tissue if reflected in the image quality in this example.

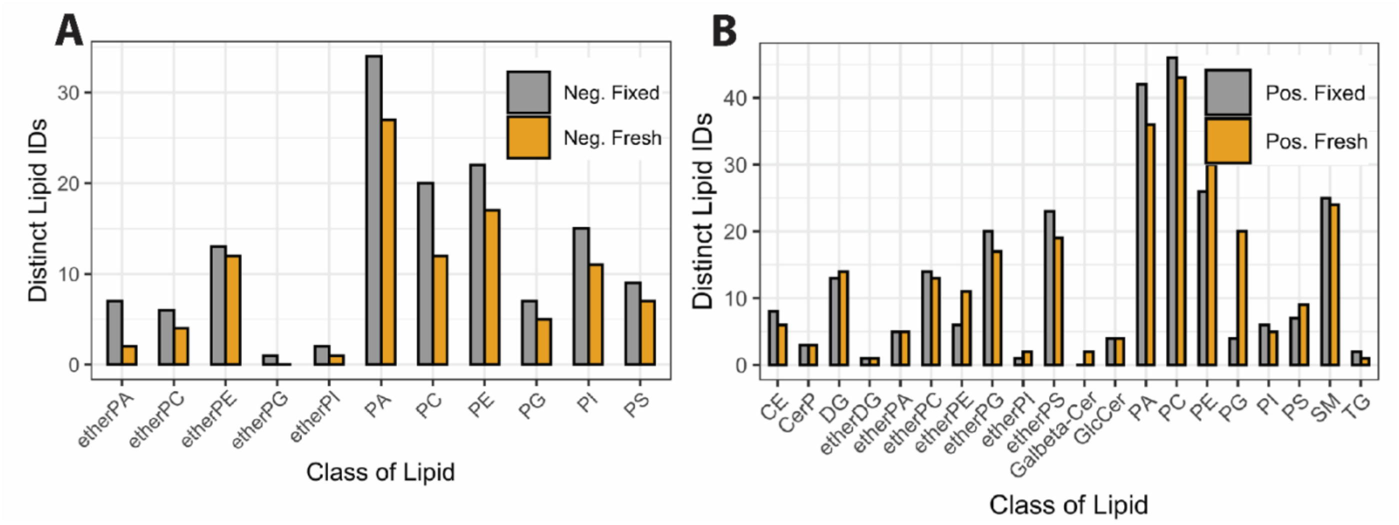


**Supplement Figure 5:** Panel A and B show the differences in lipid composition from fixed (gray bars) and fresh-frozen (yellow bars) tissues analyzed using MALDI-IMS displaying multiple lipid classes observed in negative (Panel A) and positive (Panel B) ion mode analysis.

**Supplementary Figure 6 A**: Molecular ion spectra and fragmentation patterns in negative ionization mode of lipids from fixed tissue using LC-ESI-MS/MS. Calculated monoisotopic mass **A1.** 701.5121; PA (18:1_18:0); ppm error: 1.0, **A2.** 772.5281; PE(O-18:1_22:6); ppm error: 0.6 and **A3.** 1473.981; CL (18:2_20:4_18:2_18:2); ppm error: 1.3. **B**: Molecular ion spectra and fragmentation pattern in negative ionization mode of fresh-frozen tissue using LC-ESI-MS/MS. Calculated monoisotopic mass **B1.** 700.5281; **(a)** PE(O-16:0_18:2) and **(b)** PE(P-18:1_16:0) ; ppm error: 0.1, **B2.** 722.5125; **(a)** PE(O-16:0_20:5) and **(b)** PE(P-16:0_20:4); ppm error: 0.8, **B3.** 750.5443; **(a)** PE(O-16:0_22:5) and **(b)** PE(P-18:0_20:4); ppm error: 0.3. **C:** Molecular ion spectra and fragmentation pattern in positive ionization mode of fixed tissue using LC-ESI-MS/MS. Calculated monoisotopic mass **C1.** [M+H]+ 786.6013; PC(36:2) ; ppm error:2.6, **C2.** [M+H]+ 788.6169; PC(36:1); ppm error:0.6 , **C3.** [M+H]+ 814.6326; PC(38:2); ppm error: 5.6. **D;** Molecular ion spectra and fragmentation pattern in positive ionization mode of fixed tissue using LC-ESI-MS/MS. Calculated monoisotopic mass **D1.** [M+H]+ 734.5694; PC(32:0); ppm error: 1.9, **D2.** [M+H]+ 760.5851; PC(34:1); ppm error: 2.2, **D3.** [M+H]+ 786.6007; PC(36:2); ppm error: 1.9.

**A1**

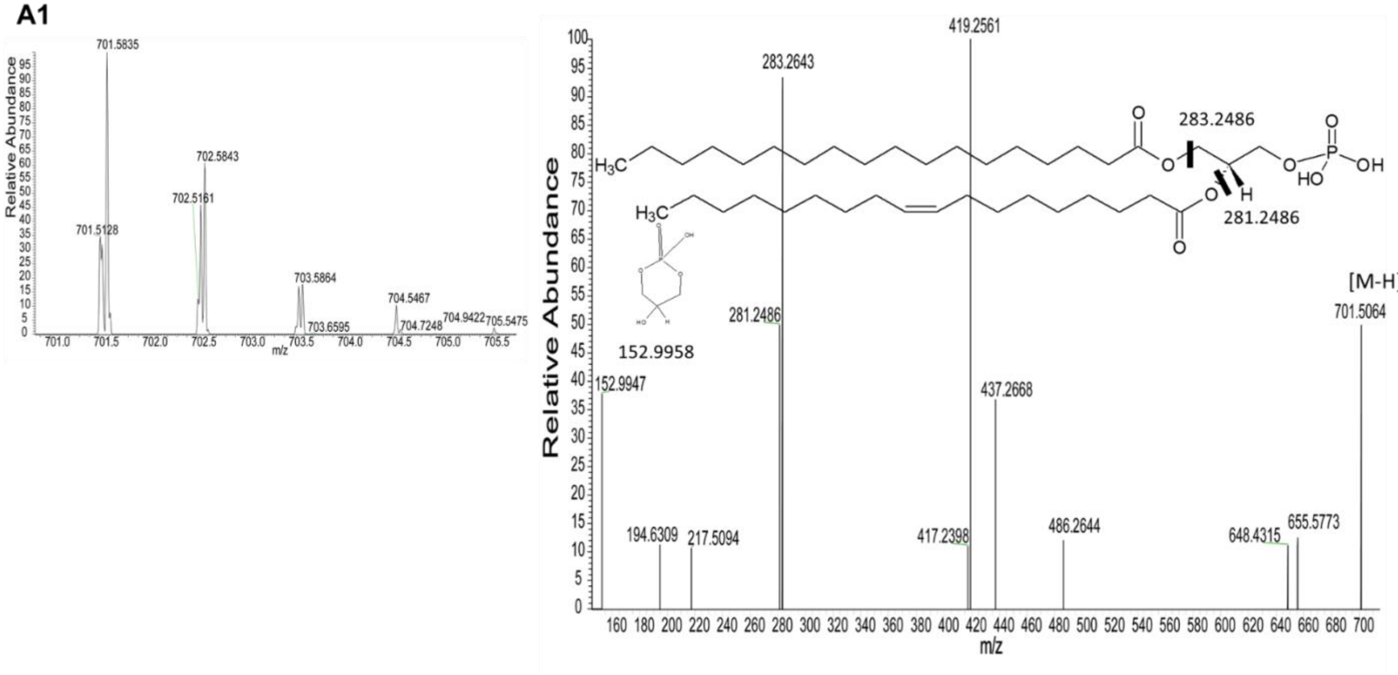

**A2**

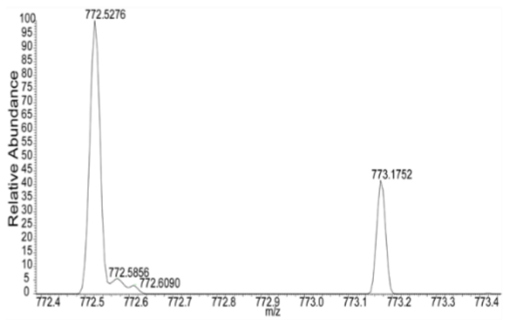
772.5276
772.5856
772.6090
773.1752
Relative Abundance
m/z

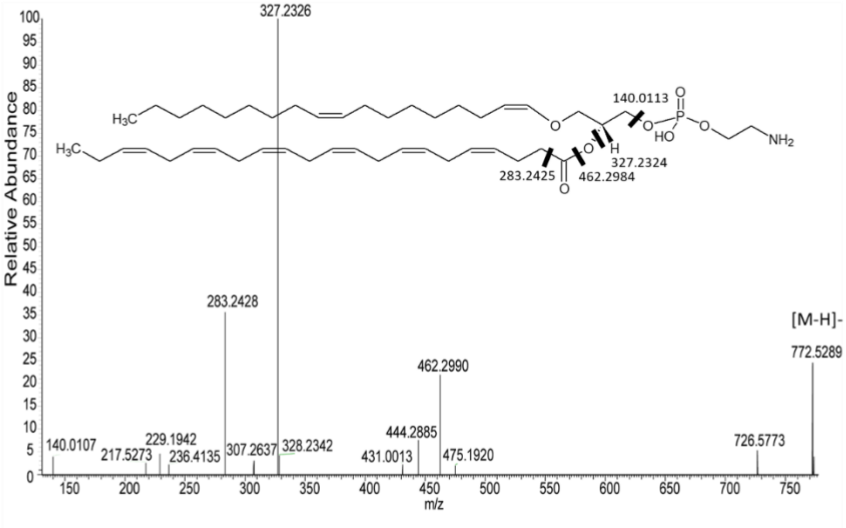
327.2326
140.0113
327.2324
283.2425
462.2984
283.2428
[M-H]-
772.5289
462.2990
444.2885
726.5773
140.0107
217.5273
229.1942
236.4135
307.2637
328.2342
431.0013
475.1920
Relative Abundance
m/z

**A3**

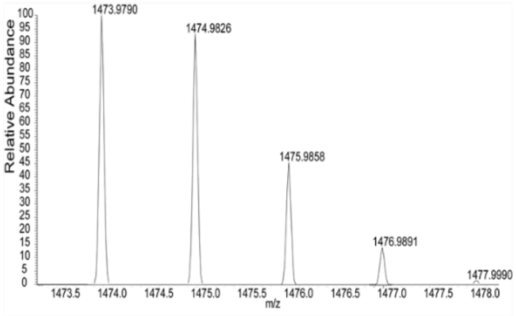
1473.9790
1474.9826
1475.9858
1476.9891
1477.9990
Relative Abundance
m/z

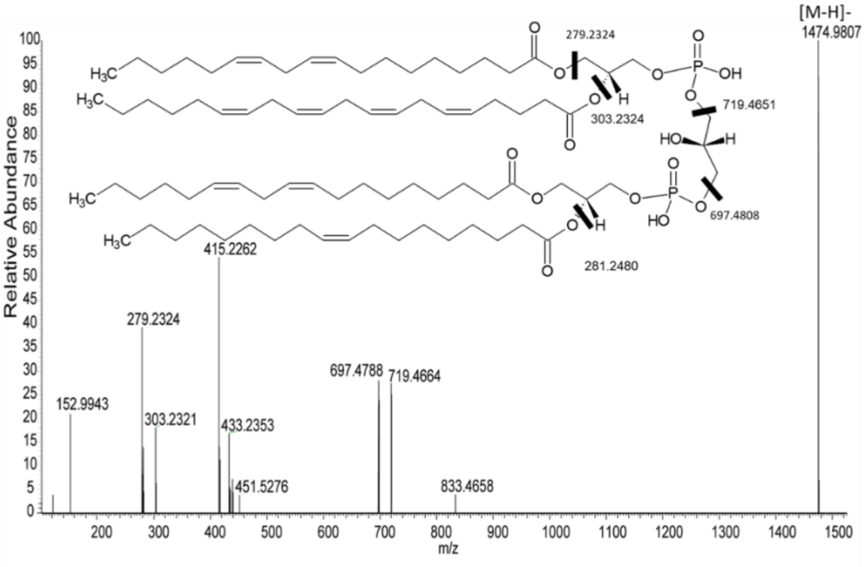
[M-H]-
1474.9807
279.2324
303.2324
719.4651
697.4808
281.2480
415.2262
279.2324
697.4788
719.4664
152.9943
303.2321
433.2353
451.5276
833.4658
Relative Abundance
m/z

**B1**

**(a)**

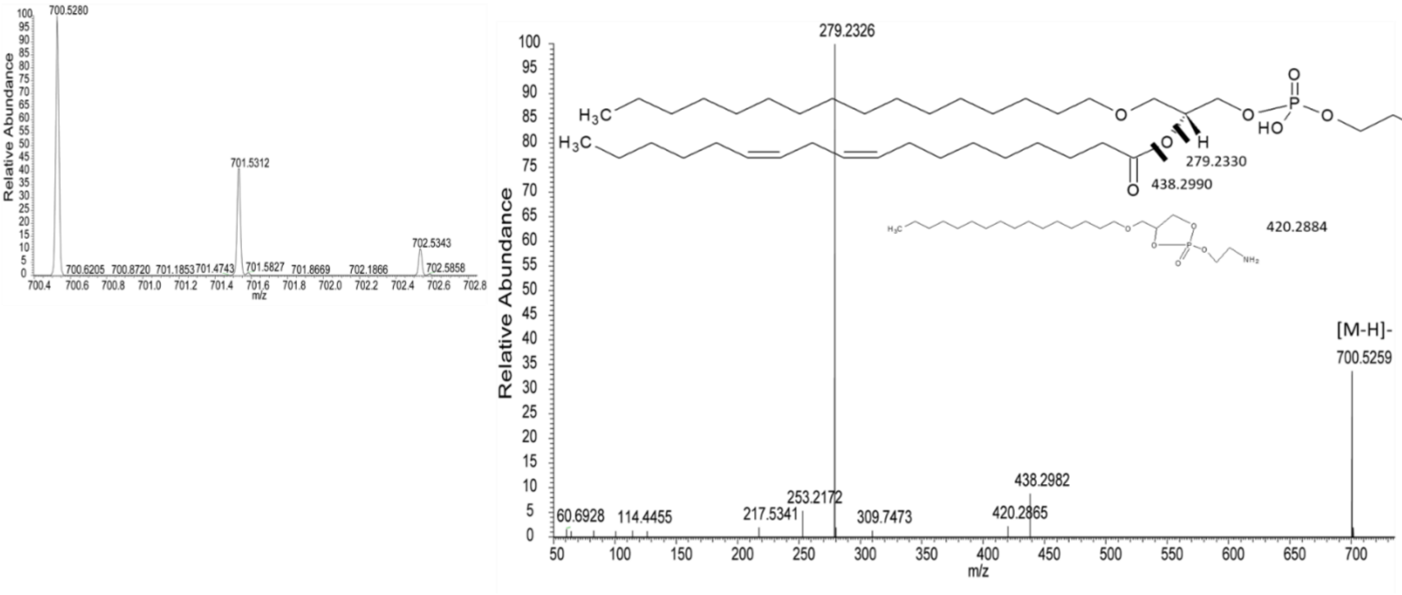
700.5280
701.5312
702.5343
700.6205
700.8720
701.1853
701.4743
701.5827
701.8669
702.1866
702.5858
Relative Abundance
m/z
279.2326
279.2330
438.2990
420.2884
[M-H]-
700.5259
438.2982
253.2172
217.5341
309.7473
420.2865
60.6928
114.4455
$H_3C$
HO
$NH_2$

**B1**

**(b)**

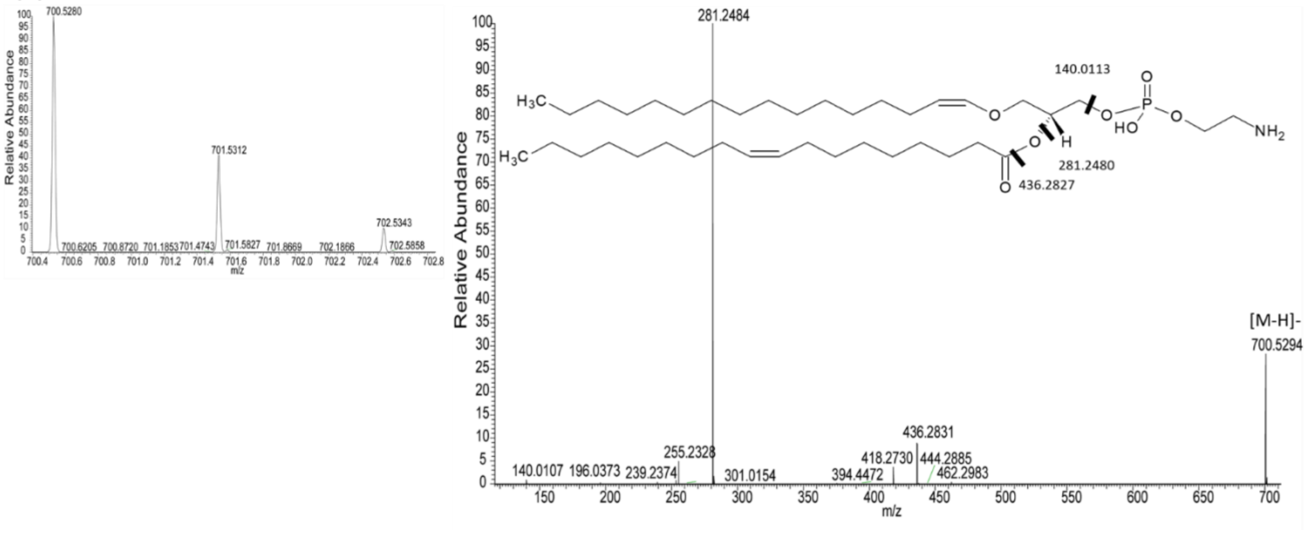
700.5280
701.5312
702.5343
700.6205
700.8720
701.1853
701.4743
701.5827
701.8669
702.1866
702.5858
Relative Abundance
m/z
281.2484
140.0113
281.2480
436.2827
[M-H]-
700.5294
436.2831
255.2328
418.2730
444.2885
462.2983
140.0107
196.0373
239.2374
301.0154
394.4472
$H_3C$
HO
$NH_2$

**B2**

**(a)**

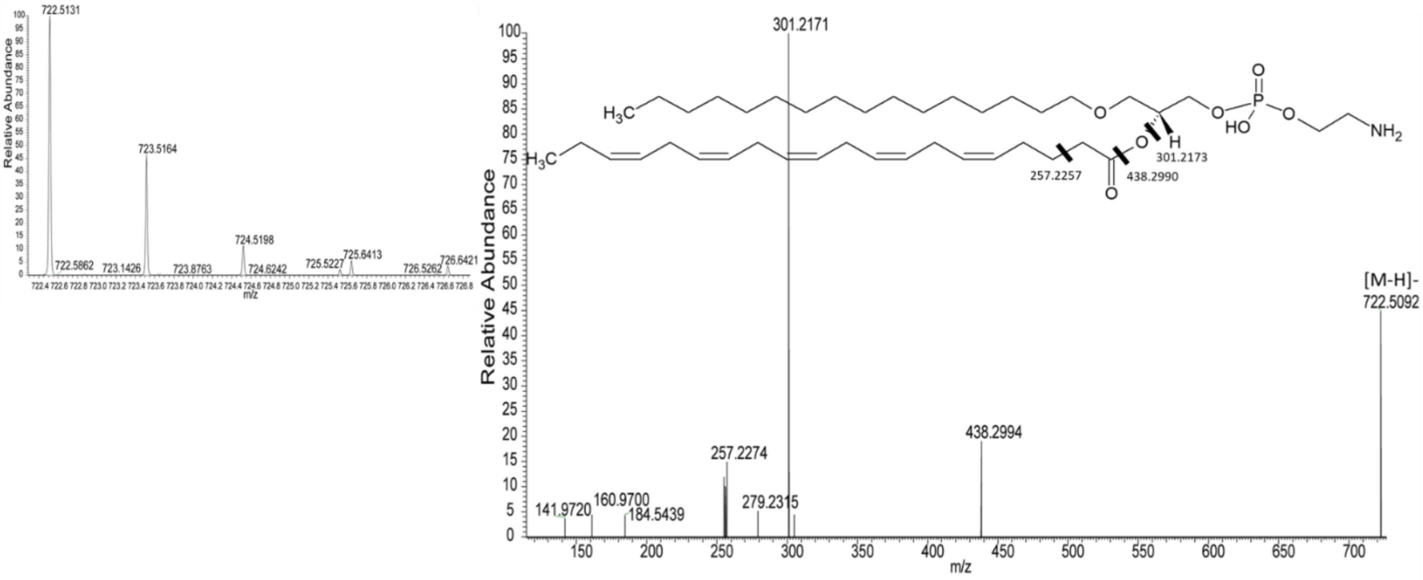
722.5131
723.5164
724.5198
725.6413
726.6421
Relative Abundance
m/z
301.2171
$H_3C$
$NH_2$
HO
257.2257
438.2990
301.2173
[M-H]-
722.5092
438.2994
257.2274
141.9720
160.9700
184.5439
279.2315

**B2**

**(b)**

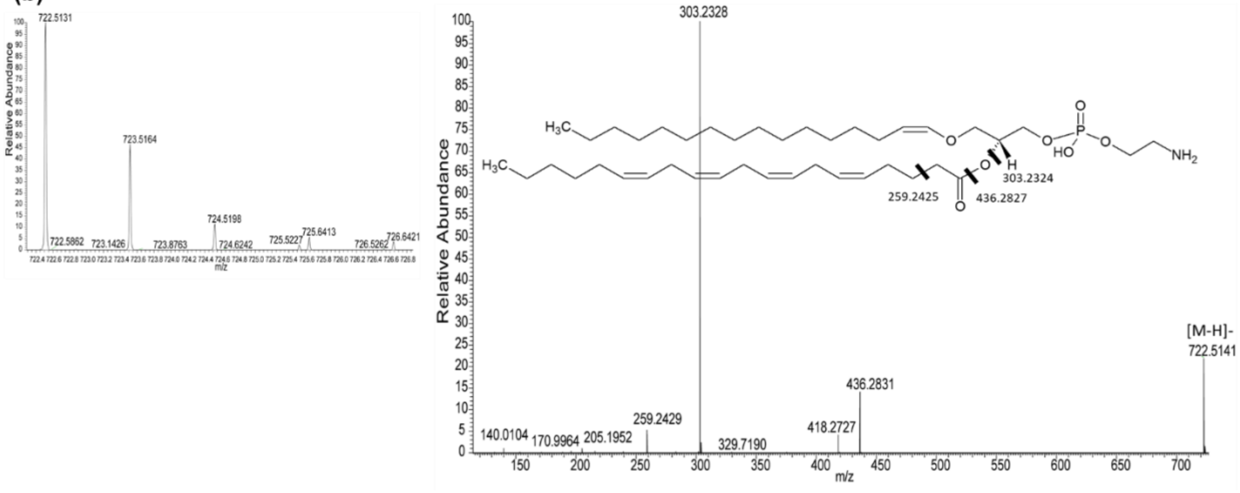
722.5131
723.5164
724.5198
725.6413
726.6421
Relative Abundance
m/z
303.2328
$H_3C$
$NH_2$
HO
259.2425
436.2827
303.2324
[M-H]-
722.5141
436.2831
259.2429
140.0104
170.9964
205.1952
329.7190
418.2727

**B3**

**(a)**

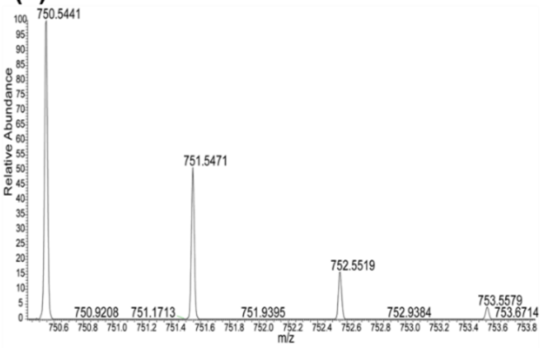

750.5441
751.5471
752.5519
750.9208
751.1713
751.9395
752.9384
753.5579
753.6714
Relative Abundance
m/z


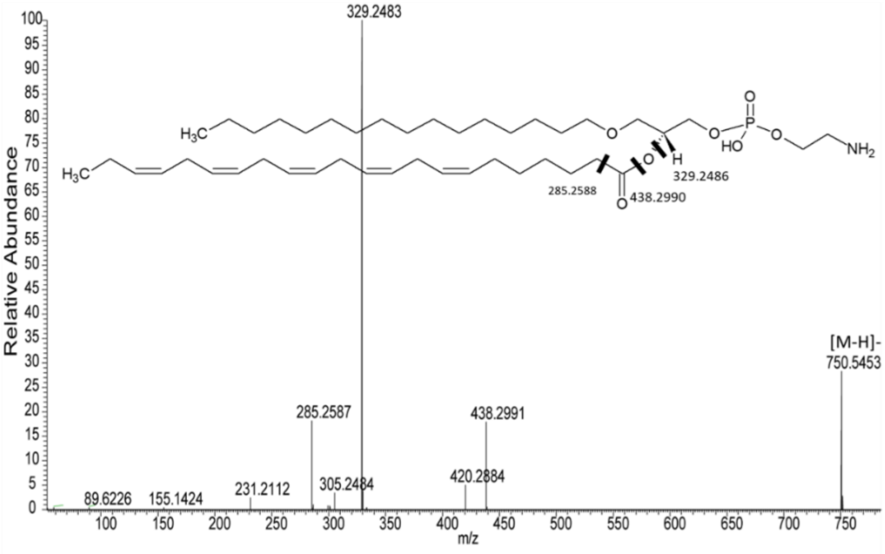

329.2483
285.2588
438.2990
329.2486
[M-H]-
750.5453
285.2587
438.2991
420.2884
305.2484
231.2112
155.1424
89.6226
Relative Abundance
m/z


**B3**

**(b)**

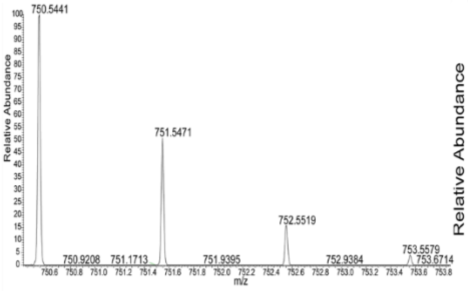

750.5441
751.5471
752.5519
750.9208
751.1713
751.9395
752.9384
753.5579
753.6714
Relative Abundance
m/z


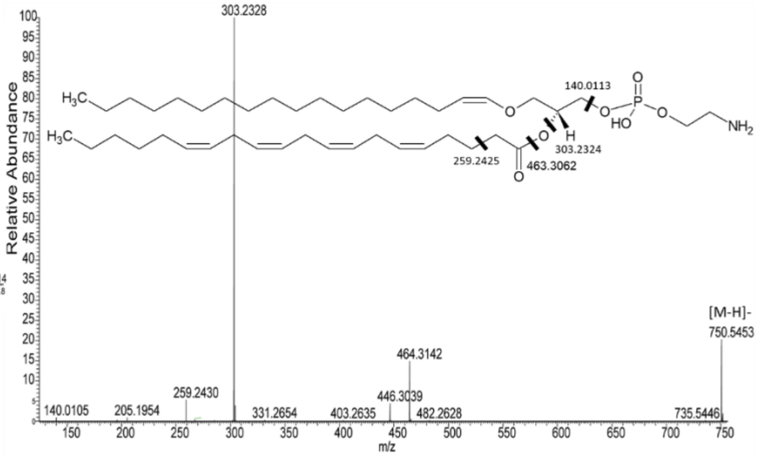

303.2328
140.0113
303.2324
259.2425
463.3062
[M-H]-
750.5453
464.3142
446.3039
482.2628
403.2635
331.2654
259.2430
205.1954
140.0105
735.5446
Relative Abundance
m/z

**C1**

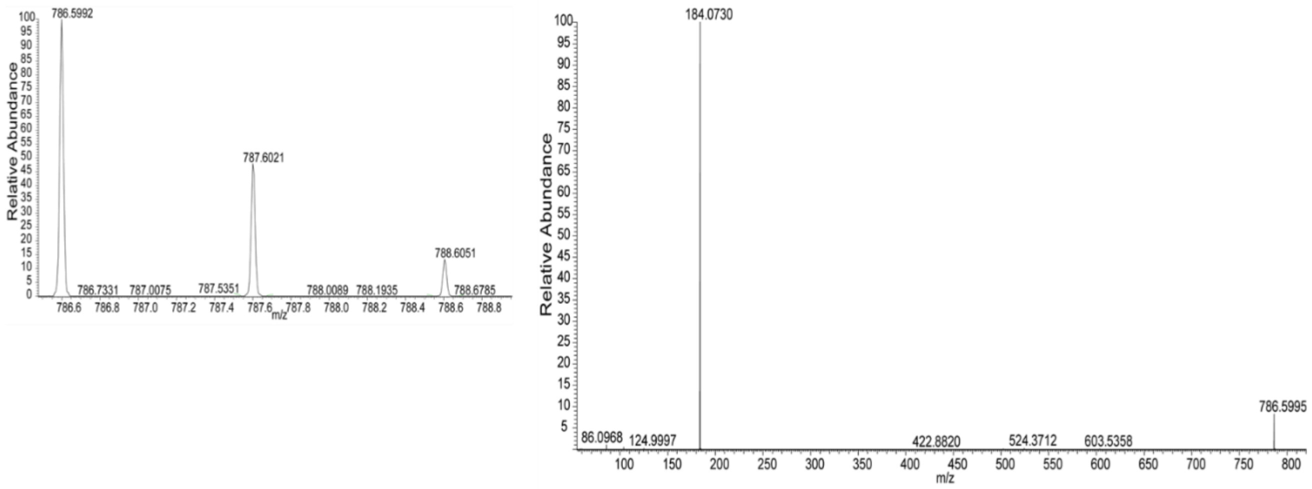
786.5992
787.6021
788.6051
786.7331
787.0075
787.5351
788.0089
788.1935
788.6785
Relative Abundance
m/z
184.0730
86.0968
124.9997
422.8820
524.3712
603.5358
786.5995
Relative Abundance
m/z

**C2**

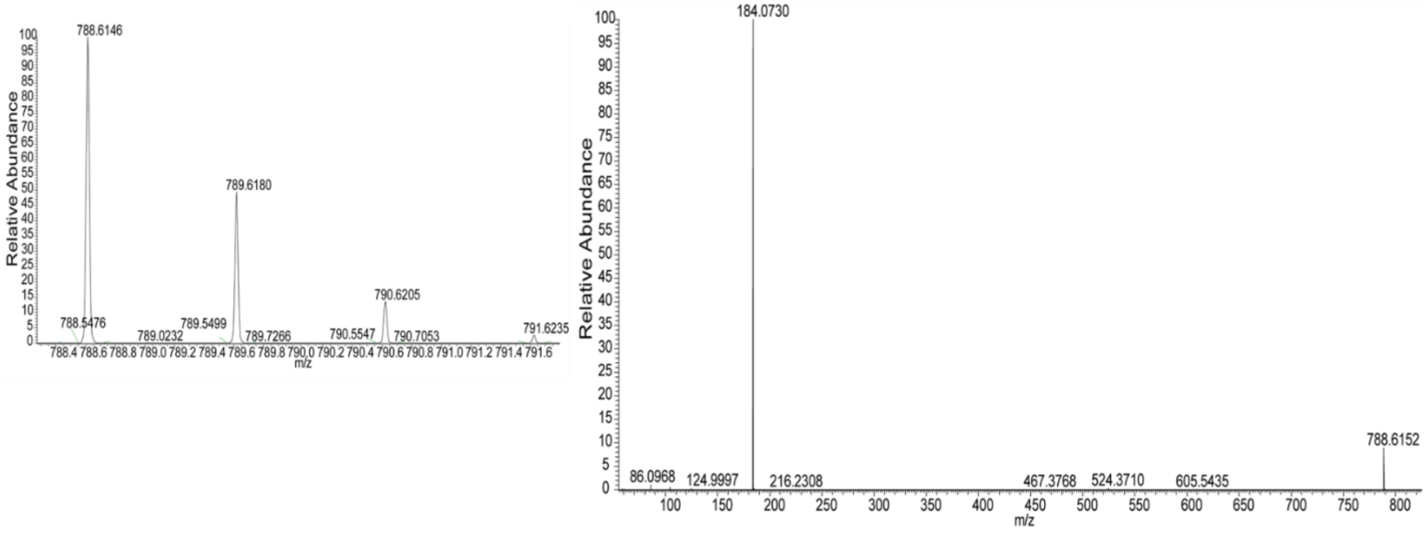
788.6146
789.6180
790.6205
788.5476
789.0232
789.5499
789.7266
790.5547
790.7053
791.6235
Relative Abundance
m/z
184.0730
86.0968
124.9997
216.2308
467.3768
524.3710
605.5435
788.6152
Relative Abundance
m/z

**C3**

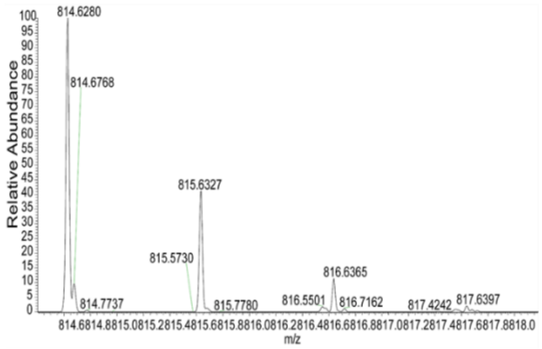

814.6280
814.6768
815.6327
815.5730
816.6365
814.7737
815.7780
816.5501
816.7162
817.4242
817.6397
Relative Abundance
m/z


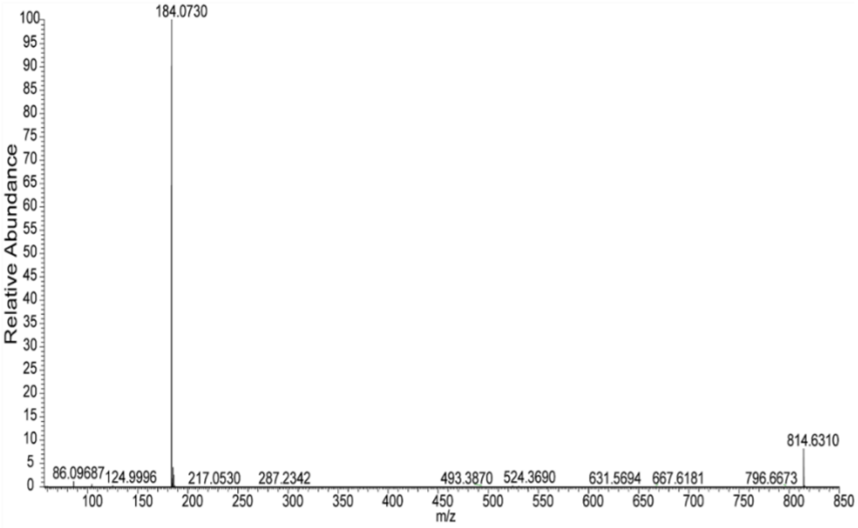

184.0730
814.6310
86.09687
124.9996
217.0530
287.2342
493.3870
524.3690
631.5694
667.6181
796.6673
Relative Abundance
m/z

**D1**

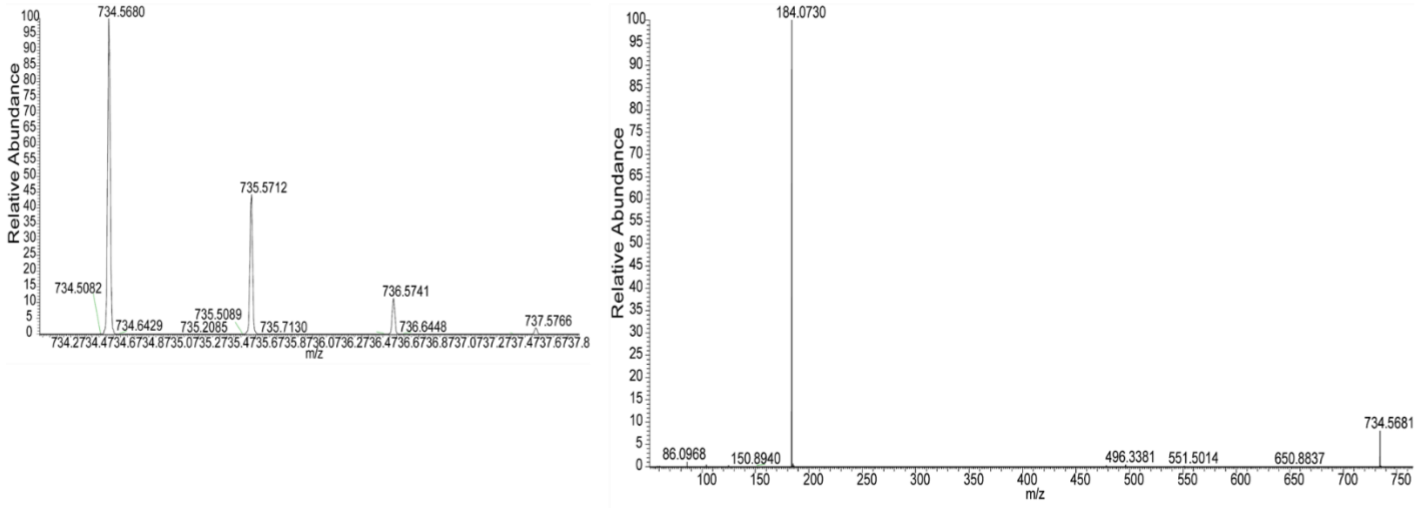
734.5680
734.5082
734.6429
735.2085
735.5089
735.5712
735.7130
736.5741
736.6448
737.5766
Relative Abundance
m/z
184.0730
86.0968
150.8940
496.3381
551.5014
650.8837
734.5681

**D2**

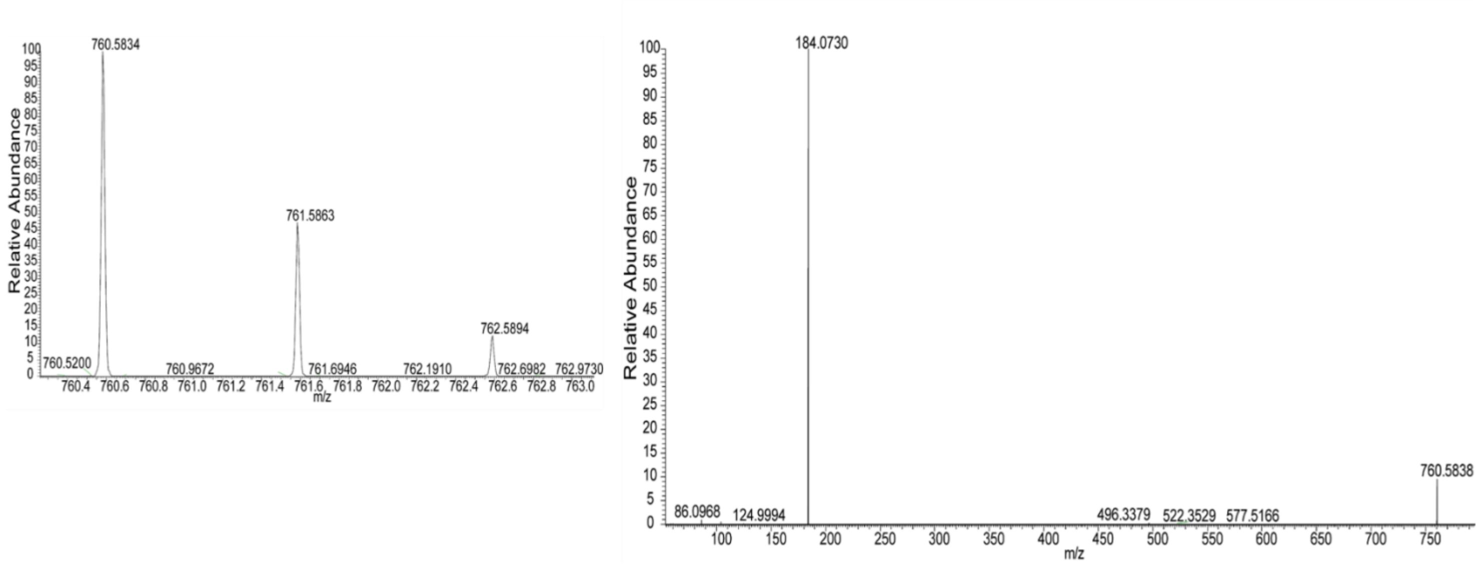
760.5834
760.5200
760.9672
761.5863
761.6946
762.1910
762.5894
762.6982
762.9730
Relative Abundance
m/z
184.0730
86.0968
124.9994
496.3379
522.3529
577.5166
760.5838

**D3**

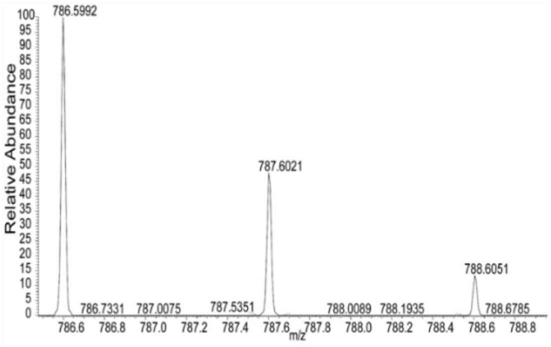
786.5992
787.6021
788.6051
786.7331
787.0075
787.5351
788.0089
788.1935
788.6785
Relative Abundance
m/z

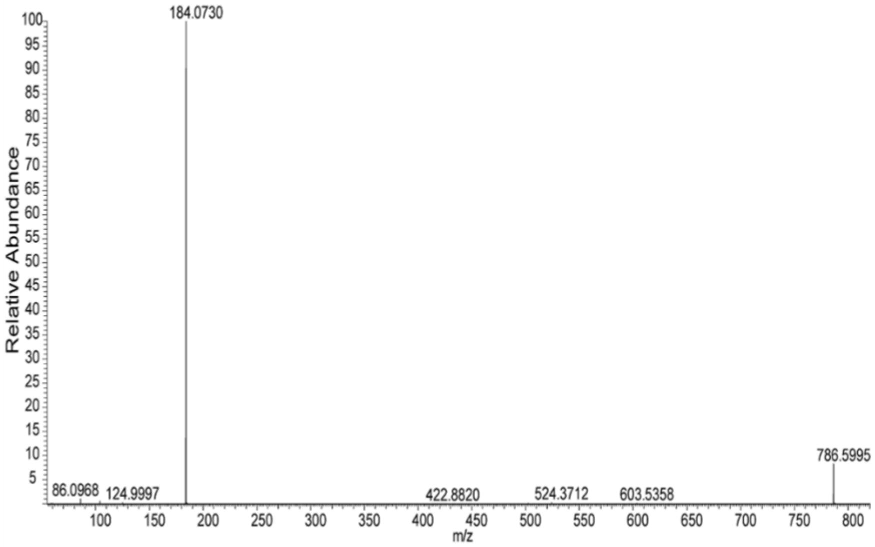
184.0730
786.5995
86.0968
124.9997
422.8820
524.3712
603.5358
Relative Abundance
m/z

**a)**

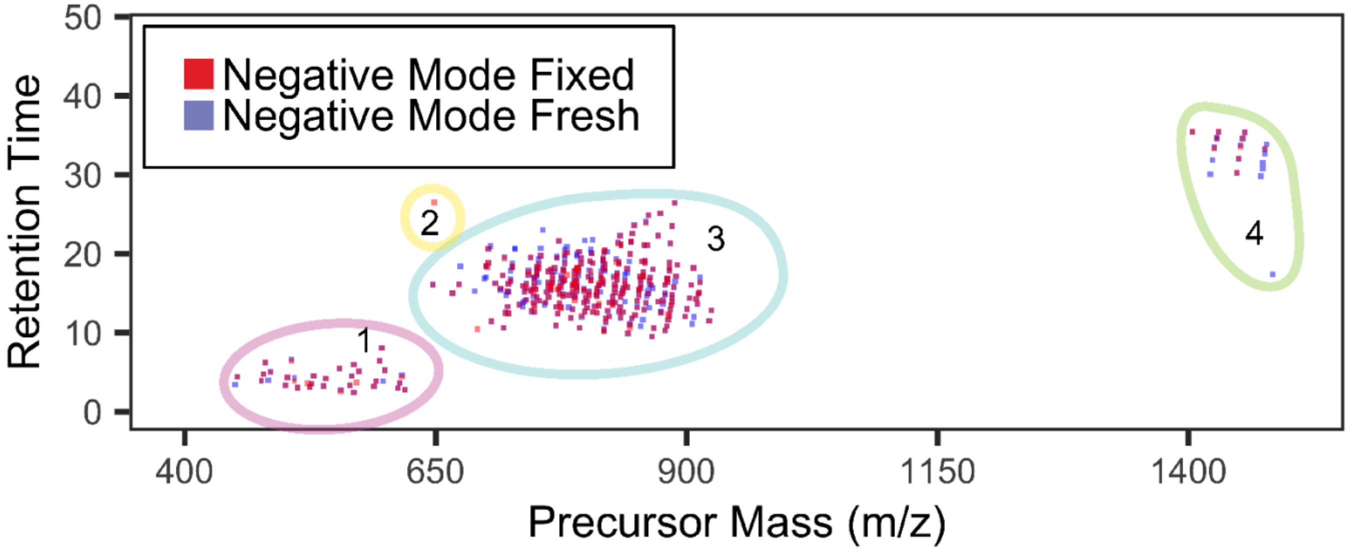


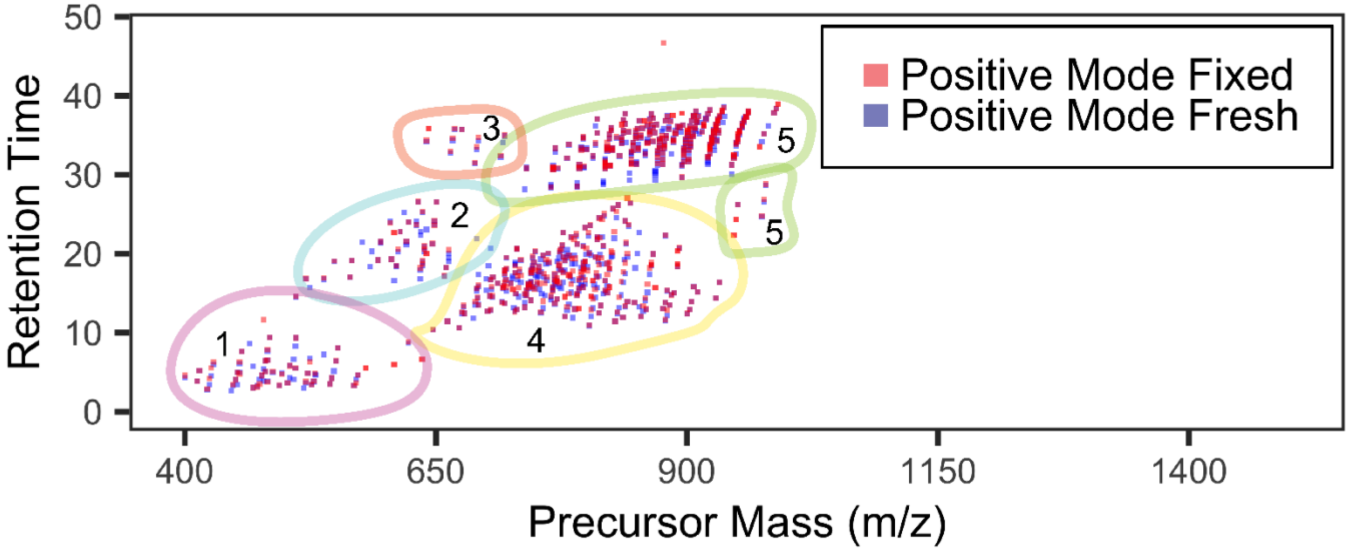


**b)**

**Supplementary Figure 7:** LC-MS/MS data of fresh-frozen (blue boxes) vs fixed (red boxes) human donor retina **(biological replicate 2)**. M/z versus retention time summary of **(a)** negative ion mode data and m/z versus retention time summary of **(b)** positive ion mode data where lipid classes.

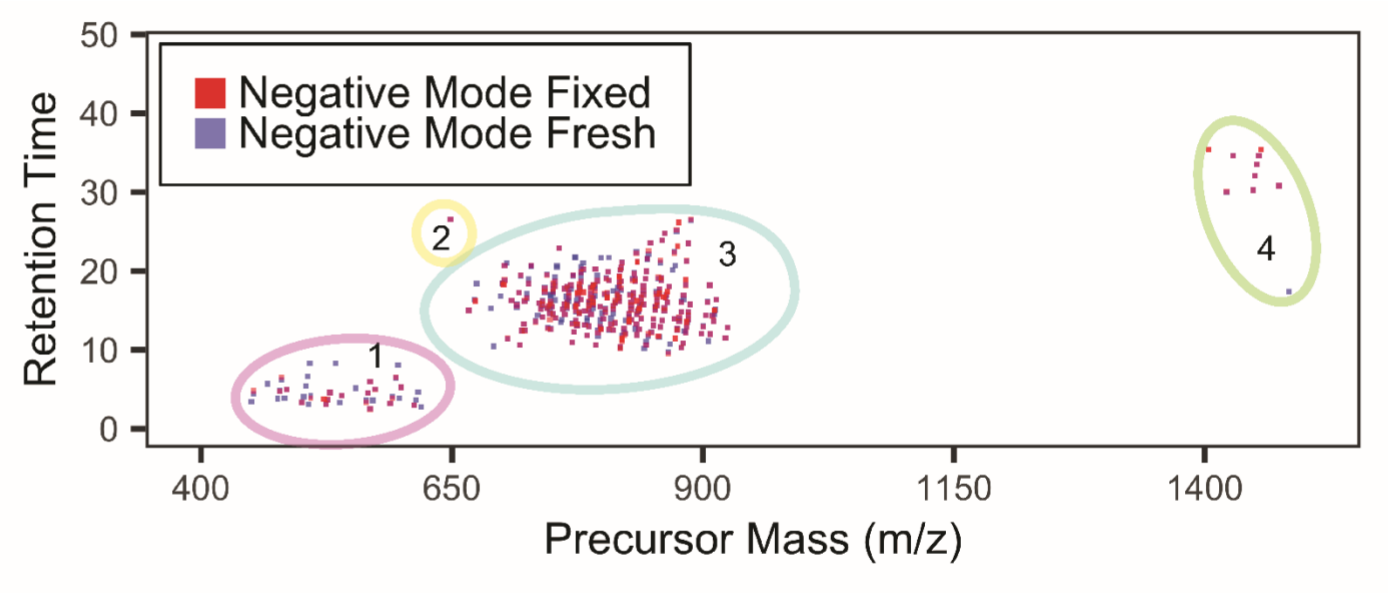


a)

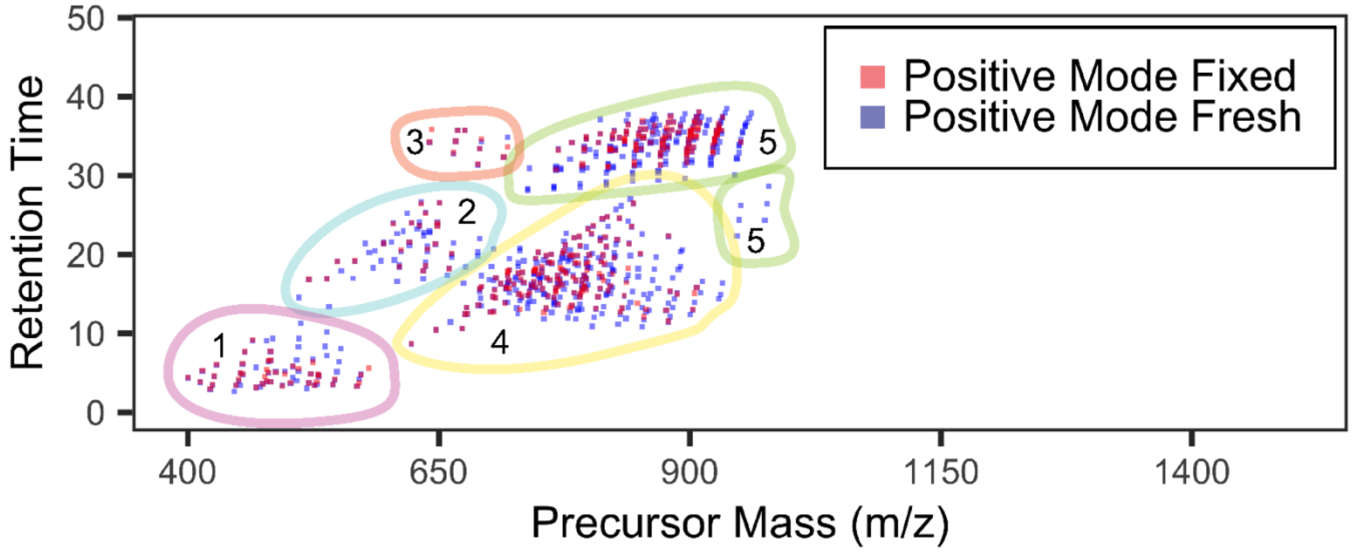


b)

**Supplementary Figure 8:** LC-MS/MS data of fresh-frozen (blue boxes) vs fixed (red boxes) human donor retina **(biological replicate 3)**. M/z versus retention time summary of **(a)** negative ion mode data and m/z versus retention time summary of **(b)** positive ion mode data where lipid classes.

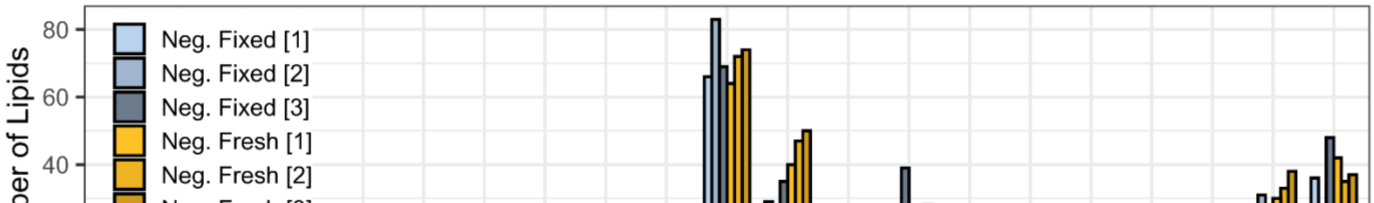

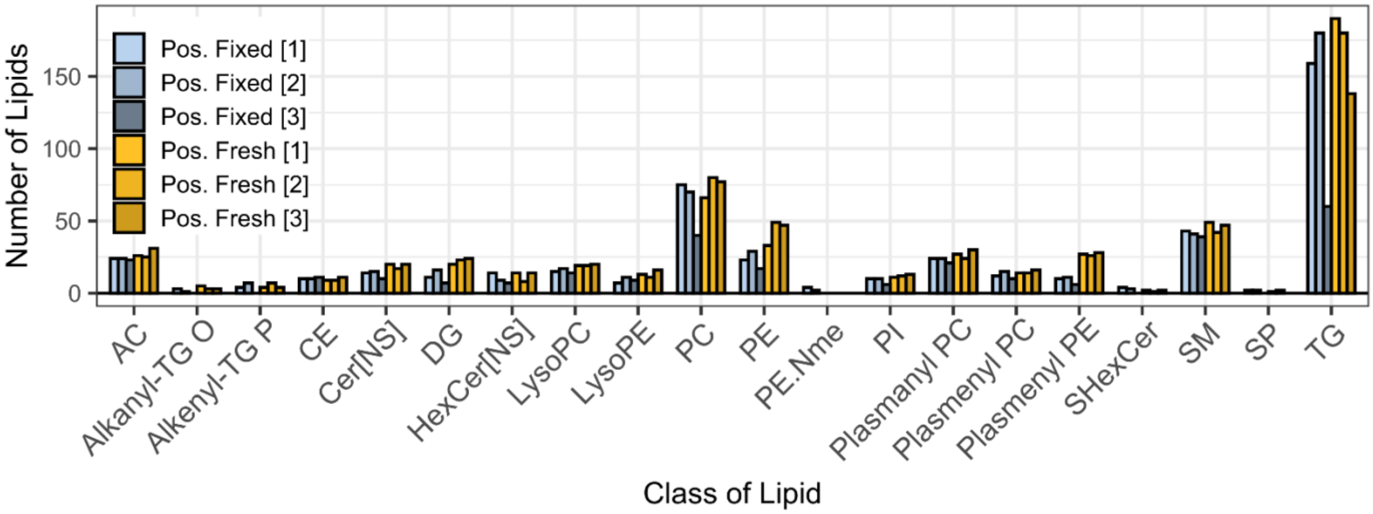


**Supplementary Figure 9:** Lipid composition identified (MS2 confirmations) in fresh-frozen and fixed human retina from LC-MS/MS using LipiDex.
